# VRoot: An XR-Based Application for Manual Root System Architecture Reconstruction

**DOI:** 10.1101/2024.06.13.598253

**Authors:** Dirk N. Baker, Tobias Selzner, Jens Henrik Göbbert, Hanno Scharr, Morris Riedel, Ebba Þóra Hvannberg, Andrea Schnepf, Daniel Zielasko

## Abstract

This article describes an immersive extended reality reconstruction tool for root system architectures from 3D volumetric scans of soil columns. We have conducted a laboratory user study to assess the performance of new users with our software in comparison to classical and established desktop software. We utilize a functional-structural plant model to derive a synthetic root architecture that serves as objective quantification for the root system architecture reconstruction. Additionally, we have collected quantitative feedback on our software in the form of standardized questionnaires. This work provides an overview of the extended reality software and the advantage of using immersive techniques for 3D data extraction in plant science. Through our formal study, we further provide a quantification of manual root system reconstruction accuracy. We observe an increase in root system architecture reconstruction accuracy (*F*_1_) compared to state-of-the-art desktop software and a more robust extraction quality.

## 1 Introduction

Understanding root systems is an underappreciated part of the process of sustainable agriculture [1, 2]. Historically, it was not possible to access root systems except by using difficult excavation processes. However, more recent research highlights that an analysis of the spatial configuration of roots, i.e., the root system architecture (RSA), is a critical aspect of understanding plant response [3]. Notably, there is a large variance in root traits [4], which impacts functional aspects of plant behavior. The root traits of a single plant can be measured from its RSA. A digitized version of the RSA yields a full description of the functional and structural traits of the root system. Functional-Structural Plant Models (FSPMs) are coupled simulations of plant structure and functional traits. RSAs can be used as boundaries in these functional simulations to provide insight into plant performance that cannot be measured directly. FSPMs bridge gaps between measurable indicators that might not directly correlate unless the plant is viewed as a continuum model [5].

Ideally non-destructive observations of RSAs are used in experiments to allow repeated measurements, such as is possible using rhizotrons for statistical descriptions of roots [6]. A full RSA reconstruction in the face of partial root obstruction or destructive measurements is challenging. Non-invasive 3D imaging methods, like Magnetic Resonance Imaging (MRI), can assist by giving a more complete insight into the root system architecture [7]. 3D imaging techniques do not require direct intervention into the plant growth, and they do not require the introduction of transparent obstructions. While the plant growth in a soil column is more restrictive than in a field, key insights can be gained from an in-depth analysis of 3D imaging data, such as the progression of diseases in the plant [8]. Another key aspect in the dissemination of information from plant image data is the potential to gain insight through root modeling, as both in-silico experiments [9] as well as quantification of the continuous processes between soil, plant and atmosphere can provide valuable insights [5].

The extraction of RSAa is more challenging and depends on the quality of the image data. Most approaches to extract RSAs from 3D image data result in tree-like or centerline structures that describe the morphology of the root systems. Fully automatic approaches fall into the categories of topological analysis [10] or optimization-based approaches [7]. There are also semi-automatic approaches, that require user interaction for key aspects, or to correct the automatic propositions. For a fully automatic approach to function, the globally optimal solution to the extraction problem must reflect the correct RSA. This is not necessarily the case, as in some measurements, artifacts due to soil composition and soil water content can lead to a difference in measured and actual morphology [11].

Manual approaches are a way of dealing with challenging image data properties, as expert knowledge can be required to completely extract the RSA to a degree that it is usable in further analysis [12]. Past approaches typically aimed at solving this challenge through guided optimization [11] or semi-automatic correction [7]. To assist with manual RSA reconstruction, Virtual Reality (VR) software has been developed, which was used to model RSA functional properties [13]. However, usability of these systems was limited as they were less portable than current VR hardware, which have improved in accessibility and display properties. The use of modern Extended Reality (XR) or VR hardware and software has the potential to increase the space of experimental conditions that are usable in combination with 3D RSA extraction techniques, thus closing the gap between data analysis requirements and realistic experimental conditions. Within VR, RSAs can be visualized directly in a more intuitive embedding, as the 3D displays will be able to accommodate and visualize depth as well as spatial configuration. VR is a promising tool, as it has been shown to improve extraction quality for similar tasks in other disciplines, such as neural imaging [14]. To improve our workflows of manual RSA reconstruction, and to investigate the applicability of VR in these workflows, we have developed VRoot, a VR application that assists in the reconstruction of RSAs by visualizing the 3D volume and providing intuitive toolsets for the reconstruction and adaption of RSAs.

We have built VRoot with the toolsets needed for exact and expedient RSA reconstruction. There are many tasks for which VR improves the quality of task completion in comparison to desktop applications. In this work, we investigate two research questions: Does VRoot improve the data extraction workflow for users annotating 3D MRIs? Furthermore, is the RSA reconstruction using VRoot more exact than using classical methods?

To answer these questions, we have conducted a laboratory user study with participants onsite. With an in-silico 3D root image, we have tasked participants with extracting the root system using VRoot and NMRooting, a state-of-the-art desktop RSA extraction and analysis application [7]. With the resulting RSA reconstructions, we have quantified key traits of the root system, and more importantly, the accuracy of the extraction in comparison to the original RSA.

This work makes several key contributions to 3D plant phenomics. First, we present a VR software to interactively reconstruct root systems from 3D imaging techniques. We quantify the user-based errors and reconstruction artifacts in a laboratory user study. Lastly, we evaluate the use of VRoot based on user feedback obtained through controlled questionnaires.

In the remainder of this section, we highlight the background on RSA reconstruction and previous approaches, particularly motivating instances in which VR is a useful tool to analyze 3D data that cannot easily be annotated automatically. In Sec. 2, we describe VRoot as well as the setup of our laboratory user study. Results of the study and our data analysis results are shown in Sec. 3, while we discuss the findings and implications as well as future directions in Sec. 4.

### 1.1 RSA Reconstruction from MRI

3D imaging techniques are methods that provide insight into the soil/root volume without the need for excavation or rhizotrons. This work focuses on the use of MRI techniques to assess the root system as there is a potential for a highly detailed extracted root architecture, depending on the image quality. 3D imaging provides more precise insight into the RSA since it does not alter the spatial arrangement and can be used to assess both morphology and topology. We refer to Haber-Pohlmeier et al. [15] for more information about the MRI systems and setups we rely on.

Fig. 1.A shows a sketch of the setup. Plant containers are typically plastic containers with a single plant growing in sieved soil. We are aiming to measure and extract RSAs from a large variety of soils and water contents. The resulting soil volume is generally a slice based 3D volume. This volume might require stitching, depending on the explicit setup of the MRI scanner. For the MRI reconstruction in general, a common approach is to quantify traits from the soil volume, as done by Dusschoten et al. [7]. There are multiple factors adding to the challenge of extracting the RSA from MRI measurements. Commonly, the soil water content together with the soil type (such as either loam or sandy loam) impacts the overall signal-to-noise ratio [16, 17]. Furthermore, ferromagnetic particles within the soil might impact the measurement quality and might even disrupt root signal continuity, resulting in gaps in the root system [11, 12].

**Fig. 1:**
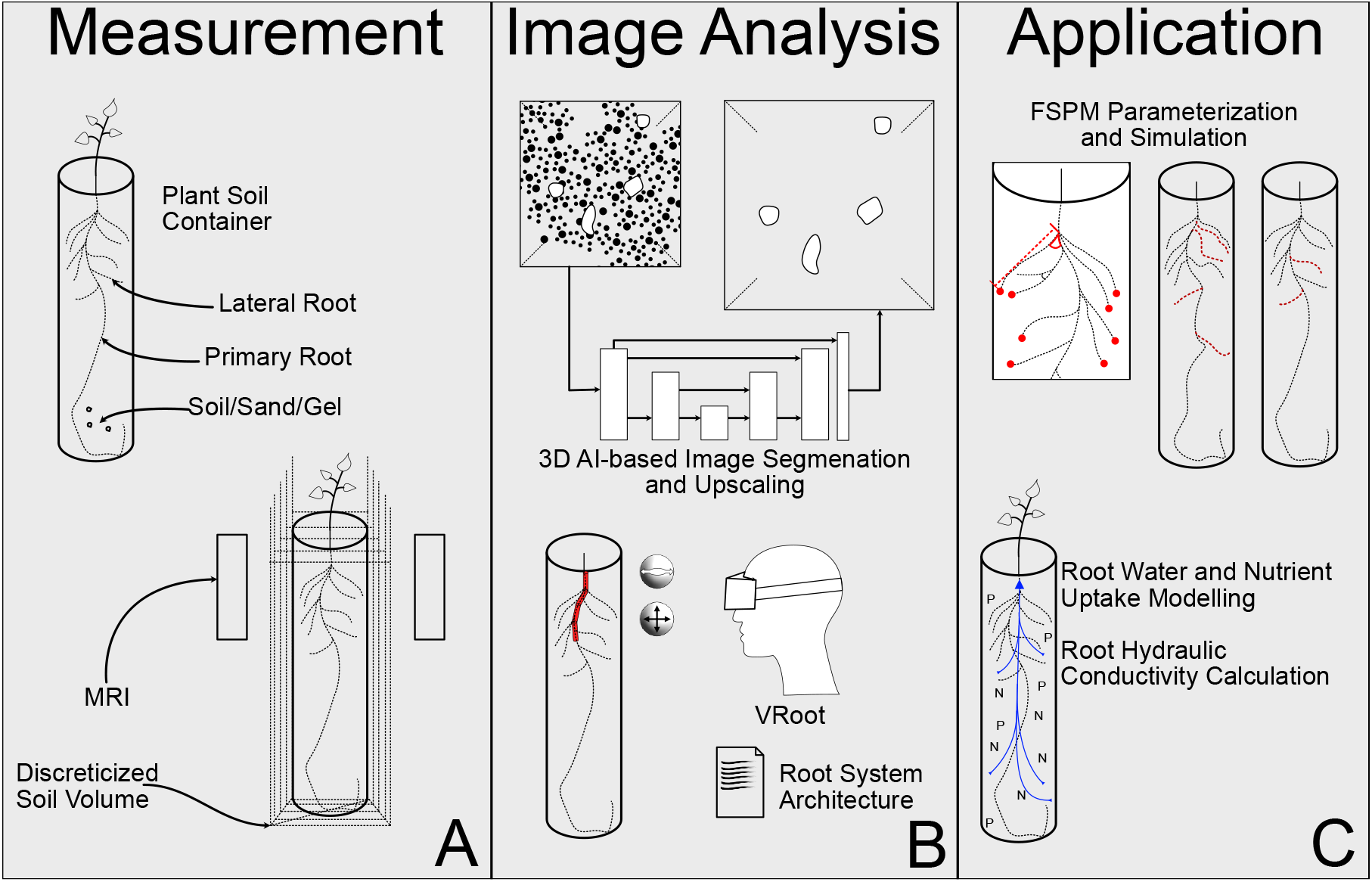
A: Plant containers are non-magnetic tubes with sieved soil. These are scanned using an MRI scanner, resulting in a voxelized soil volume. B: If the data is too noisy, 3D-U-Net segmentation can be employed in addition to the explorative functions of VRoot. C: The resulting RSAs can be used for FSPM calibration or functional simulation boundary.

To combat the potentially low quality of 3D image data, there has been notable advancement in segmentation procedures that assist with the pre-processing of soil imaging. Segmentation is generally a voxel-based mapping that labels a part of the image as either foreground or background. Elliot et al. [18] have investigated the use of different thresholding techniques. This approach is especially helpful, as it is easy to compute, primitively parallel, and is a good first indicator on image data quality. Thresholding and/or isosurface extraction is an expedient method of displaying 3D image data in VR and is used as explorative feature in VRoot. More complex segmentation approaches have been developed by Douarre et al. [19] for X-ray tomography. Notably, the use of deep neural networks has become very popular recently, with approaches focusing largely on segmentation, such as by Soltaninejad et al. [20], rather than other parts of the extraction pipeline. Upscaling the 3D measurements while also segmenting the image data could further improve the processing for automatic algorithms. Zhao et al. [21] and Uzman et al. [22] worked on upscaling MRI images using convolutional neural networks that have additional processing and upscaling layers. These super-resolution steps improves annotation in VR, as reported by Selzner et al. [12], who have investigated key characteristics of RSA reconstructions done with and without the assistance of super-resolution steps in the pipeline. Fig. 1.B contains an illustration of this step. Generally, network architectures vary, but the key challenge for segmentation networks is providing a full view of the root system in the soil, while removing noise from it. This reduces the visual complexity of the image data both for the benefit of automatic approaches but more impactfully for increased visual quality when viewed both on a desktop as well as in VR. Segmentation improves visualization performance by removing noisy artifacts that would otherwise contribute to an increased amount of background information that needs to be rendered by, e.g., isosurface renderer.

In terms of automated extraction approaches, most methods generally rely on sparse extraction from volumetric data. The approach developed in Van Dusschooten et al. [7] uses a voxel signal cutoff to determine what areas could include root voxels, followed by a signal-strength-weighted shortest-path algorithm to extract the root system. A similar technique was implemented by Horn et al. [11], who conducted a direct comparison to manually annotated RSAs. Their algorithm optimizes signal-based features. However, they highlight a case in which this locally results in an incorrect tracing as compared to expert annotation, because the algorithm chooses smaller (more optimal) gaps based on signal strength, while the expert annotation bridges a comparably much larger gap in the segmented image data. This kind of expert knowledge is hard to incorporate into automated algorithms in cases where the information cannot be gained otherwise, for example through repeated measurements or measurements at a later point in time when the roots have developed further. The issue of proximal roots was tackled by Zeng et al. [10] using an algorithm that simplifies the geometric topology of a structure. This approach directly uses the outline geometry of the root system respective to a specific signal cutoff value, resulting in an isosurface of MRI cells that have the same signal value. Center-line algorithms, such as topological thinning, are sensitive to features of topological significance, such as holes. Zeng et al. use a heuristic that simplifies the homotopy group of the geometry [23].

To store and analyze the extracted RSAs, both common and novel data structures have been developed. The Root System Markup Language (RSML) [24] is an extensible way of encompassing the complete plant structure. It is based on the XML standard and can be extended to other use-cases. On the other hand, multi-scale structures such as Multiscale Tree Graph [25] provide a more rigorous definition of a file format and leave less room for deviations from standardized descriptions. Notably, RSML is able to accommodate functional properties as well as definitions of shapes and a variety of organs. Polyline-based descriptions, such as files associated with the Visualization ToolKit (VTK) [26], are also prevalent, for example as an output type in applications such as NMRooting [7].

Ultimately, the VR application presented in this work provides an immersive visualization of root systems and can easily be coupled to automatic tracing algorithms through their standardized output. Our application furthermore benefits from segmentation approaches that enhance the spatial visibility of the root morphology [12]. Lastly, using our approach, one can use previously unusable image data, either because automated tracing algorithms still fail in certain cases, or because a level of precision needs to be reached that would be otherwise unobtainable with desktop software.

### 1.2 Immersive Analytics

Visual Analytics is a discipline of supporting data analysis and reasoning through visualization and graphical interfaces [27]. A subset of this field, and the collection of techniques it encompasses, is Immersive Analytics (IA), which involves the use of immersive interfaces, such as VR, which itself is a subcategory of XR. There have been a large variety of use cases for IA in science, and the VR application described in this work is another example. In their analysis of the field and its uses, Fonnet and Prié [28] evaluated the literature and the techniques used by different applications comprehensively. Notably, there are a lot of purely analytical tasks, ranging from navigation, selection, and interactive level-of-detail approaches. However, there also have been use-cases for annotation, especially sparse annotation of 3D image data. We focus on use-cases that are comparable to the use of VRoot, to provide context and an overview of instances in which IA has provided increased insight for data analysis pipelines.

The neuron tracing application developed by Usher et al. [14] use Head-Mounted Displays (HMDs) for the sparse annotation of 3D image data. Usher et al. [14] developed an application to trace neuron connections in 3D space with handheld controllers and consumer-grade VR hardware. Usher et al. show that there is a significant speedup for experts to annotate neuron traces in VR in comparison to a classical desktop application. This result has been further improved by the introduction of topological features assisting with the extraction of neuron traces as shown by McDonald et al. [29].

Immersive displays are varied, ranging from room-scale installations to small portable devices. In comparison to the ImFlip150 system used in Stingaciu et al. [13], HMDs require less space, are mobile/movable, and can be comfortably used at any office workplace. A disadvantage of HMDs in this comparison can be the loss of reference in the real world. Zielasko et al. [30] use, for immersive exploration tasks, a virtual representation of a physical desk for spatial reference to mitigate this effect, in addition to VR-sickness effects some users experience. There are systems where the own body is still visible through either pass-through view or avatar representation, such as analyzed in the context of location memory by Murcia-López and Steed [31].

The HMD used in this work, and other HMDs as well, typically are wearable low-persistence displays [32] that are tracked by either using base stations (such as the HTC Vive) or use inside-out tracking that estimates the user’s position through cameras. Users interact with the virtual environment by using trackers, which are reference points that the user wears or holds, which have 6 degrees-of-freedom (DoF) tracking of position and orientation. VR controllers contain trackers in addition to buttons that users can press for certain interactions. Interaction in VR is commonly done using interaction metaphors, which are user actions done using controllers or gestures that impact the virtual world, as the user cannot directly interact with it. These include grabbing to virtually pick up items, as is common in VR applications, but also pointing for movement outside of the restrictions of the installation or walkable space [28].

Evaluating the use of human-centered techniques, particularly in the case of immersive systems, is challenging [33]. However, there is a long history of formal analysis in human-computer interaction that we can make use of. For example, a common method of evaluation by users is the System Usability Scale (SUS) [34], which introduces the notions of usability regarding task completion. Questionnaires such as SUS have been predominantly designed for software system evaluation but can be used for immersive software, as the inherent effects are similar [35]. Evaluation metrics used in this work are described in Sec. 2.4.

### 1.3 NMRooting

NMRooting [7, 16] is a framework and application for the extraction of root system architecture by extracting the minimum-weight shortest paths with additional functionalities, such as gap closing and semi-manual extraction. We chose this application as a baseline as it is a well-established application that is also a proxy for similar applications and approaches, such as those developed by Horn et al. [11] or Zeng et al. [10]. Furthermore, NMRooting has seen regular use, including recently by Le Gall et al. [36], who used the non-destructive investigation of the RSA through NMRooting to analyze whether the root water uptake profile is an indicator for plant development.

NMrooting uses desktop user interaction metaphors such as clicking and dragging to fulfill 3D annotation. Clicking in NMRooting traces a selection ray from the viewing surface to the isosurface data, marking the first surface that it hits. Dragging is a metaphor that allows users to turn the camera around the isosurface data, as well as zoom and pan. Within NMRooting, users can alter the automatic tracing of the data set, which is more in-line with the use of applications such as TopoRoot [10] or the application developed by Horn et al. [11]. However, more fine-granular alteration to the reconstructions are possible by directly interacting with the data, yielding higher extraction accuracy in cases such as larger gaps as reported by Horn et al. [11].

## 2 Methods

### 2.1 Virtual Reality Root Tracing

We present VRoot, an application for manual RSA reconstruction and correction. To obtain the RSAs with the accuracy that we require, we need a tool that allows expedient fine-tuning. Manual annotation in VRoot is loosely based on methods developed and used by Stingaciu et al. [13] and Usher et al. [14]. A key aspect of the rendering of 3D imaging data in VR is the ability to look at the data from different perspectives while retaining the dimensionality of the data regarding their perception. We implemented the application with the key idea that different users might want to interact on different scales, in different postures [37], or with different configurations. For example, the application can be used sitting with a relatively small MRI visualization, or used standing with a fairly large visualization. While there are considerations on accuracy, as large MRIs might be easier to fit a root into, this is ultimately a matter of personal preference.

VRoot is an application that assists with very accurately tracing RSAs in 3D volumes, by providing immersive visualization and intuitive interaction. Effects that contribute to the need for manual correction include low MRI quality [12], complex root systems, or the need for very high accuracy. To this end, VRoot renders an interactive visualization of the underlying RSA by providing a node-link-based visualization of the RSML structure. This graph structure is editable at its joints, and the application provides means of selection, and tuning of the structure to accommodate a more precise extraction. In the application, the soil volume is thresholded and visualized as isosurface computed around a cutoff value that can be chosen dynamically within the application. We use the Visualization ToolKit (VTK) [26] as back-end for the visualization, running remotely, making it possible to use a large-memory server to accommodate large data sets. VRoot can be used to extract RSAs from MRIs and CTs in different resolutions and file types, including image stacks. To efficiently render the isosurface in the front-end, we simplify the surface geometry on the server by merging triangles of similar orientation, which reduces the amount of information needed to describe a similar geometric structure, before sending the geometry to the VR application.

Fig. 2 shows sample views from the user’s perspective in the application. We chose a darker environment to reduce eye strain. Users interact with two controllers, one for selecting as well as tracing, while the other controller is used for grabbing metaphors.

**Fig. 2:**
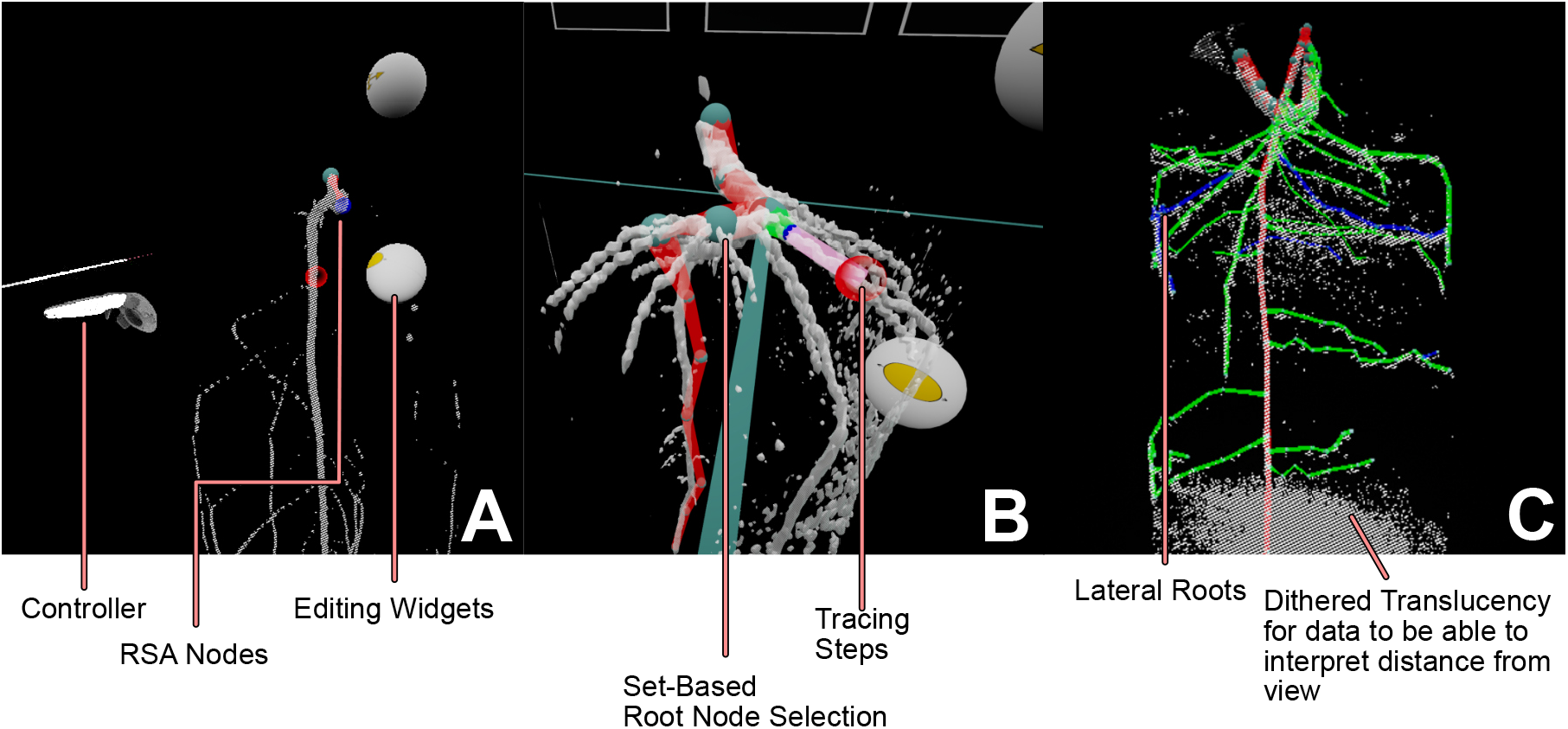
A: Two-handed use of the drawing function. New position and connection is indicated in pink. B: Selection is set-based and changes are done on all selected nodes, drawing is only possible with one. C: Subsequent root orders have high color contrast and MRI is dithered for depth-preserving translucency.

Fig. 2.A shows the basic user interaction components. Interaction with the RSA is done via the nodes. All nodes in the RSA are selectable and while tracing depends on manual user interaction, changing of properties is selection set-based, meaning that users can mark as many nodes as desired to change their properties. The widgets (grey) change the selection set’s properties, such as diameter and position. Fig. 2.B shows a snapshot of a user drawing a root, indicated by the pink interaction. This is automatically available once the number of selected nodes is exactly one, or without any tracing present. The widgets to edit node properties always work on the selection set, changing the diameter or position of the selected nodes. The volume itself is displayed as an isosurface, as seen in Fig. 2.C, whose signal cutoff value can be changed from within VR through the use of a slider widget. The full visualization of the RSA always uses the RSML topology and assigns colors to root order. We are using dithered translucency for the isosurface to avoid depth-perception issues with the surface. More general root functionality that is outside of node editing is accessible via a point-and-click menu, such as assistance tools for time series annotation or editing tools for RSA topology. Users can generally place nodes freely within the 3D environment, though it needs to be stressed that there is a degree of freedom in VR, the rotation, that is not captured by common data types, such as RSML or VTK format. To enable a more expedient workflow, many more complex tasks, including the visualization algorithms, are offioaded onto a server to allow the interactive application to run smoothly while still allowing the completion of complex tasks. For a selected data set, the system searches for the most recent tracing and displays it on top of the 3D image.

We designed the application with expert workflows in mind and have been continuously improving the workflow to assist with observations such as by Stingaciu et al. [13] and Selzner et al. [12]. However, the assessment of a system that depends on human interaction is not as straight forward as assessing the quality of an algorithm, even with a ground truth data set present. We are evaluating this in a more controlled fashion by performing a user study that is being guided by synthetic data and user questionnaires, under the restriction of using a pool of potential users that all have a similar knowledge level about the applications. We chose to evaluate untrained user performance for this reason, allowing us to focus on the relative performance of the applications without needing to balance the data for previous experience.

### 2.2 Laboratory User Study

To answer our question on whether VR annotation can outperform state-of-the-art desktop annotation for RSA reconstruction, we performed a mixed design laboratory study, assessing the applications within-subjects with the between-subject condition of water noise. We compare our software against NMRooting described above, since NMRooting is not only state-of-the-art for 3D annotation, but also is similar to other applications. The evaluation of the study is aimed to answer the question on whether the VR software yields a higher reconstruction accuracy as well as a higher usability. We map performance to reconstruction accuracy in a virtual MRI scan: Our comparison between applications and conditions relies on the assumption that how closely a participant (after a short training phase) follows the ground truth with their annotation is a direct indicator of the usefulness of the application. In this instance, the term laboratory study refers to a controlled setting in which human participants with similar starting conditions could perform tasks and evaluate the applications. Through the use of a sufficient number of individual participants, effects that are individual to certain people should be eliminated, and the overall usability of the software can be evaluated. To enable this process, the set of possible options within a single application has to be restricted, so we exclusively use “tip-to-tree” and node annotation in NMRooting and basic drawing without correction in VRoot.

The user study was designed to answer our questions and assumptions on the improvement of software and measurement quality from MRI scans. We postulate the following hypotheses on the application performance on the software level as well as on the data level.

#### Improved Workflows

We expect that the major indicators for software quality will be improved when using VR software. These indicators are an improvement (**H1**) of System Usability as well as an improvement in the subjective pragmatic performance (**H2**) of the software. These hypotheses will be tested using the participants’ evaluation using the questionnaire after task completion.

#### Extraction Accuracy

For the extracted root systems, we postulate that key relevant measures will be impacted by the use of VR. The total root length is expected to be different (**H3**), which also applies to the branching density (**H4**). We believe that the VR software will result in a higher overall accuracy (**H5**) which is further impacted by the presence of water noise (**H6**). Water noise and impact of signal-to-noise ratio have well-reported impacts on reconstruction accuracy [7, 10-12], which is why we assume that it is a significant factor in the reconstruction accuracy of participants. We expect that the total root length is closer to ground truth when using VR (**H7**) and that the VR extraction of the branching density does not differ from the branching density within the virtual MRI (**H8**).

#### 2.2.1 Tasks & Measures

The main task that participants were asked to perform is extraction of the RSA from an MRI soil column scan. This includes the extraction of the pathway of individual roots as exhibited within the MRI scan, loosely based on signal strength. Participants were asked to mitigate noise effects if present and extract a fairly simple explanation for the signal that they were shown. Furthermore, participants created a *labeled* RSA, which includes the order of the root explicitly. Participants were asked to label both primary and lateral roots as such. The tracing of the RSA resulted in each case in a full RSA, including positioning but excluding diameter. Participants were asked to provide their demographic information as well as a subjective evaluation of the software they were tasked with using. In total, participants performed the extraction task two times, with the evaluation of a questionnaire in between and at the end.

Participants were tasked with extracting a root system from an MRI scan, once using the VR application and once using NMRooting. The water noise condition spanned both data sets, meaning that independent of the order, a participant either completed the task for each application with water noise, or without. Participants were tasked with tracing a virtual MRI scan, as described in Sec. 2.3. Extraction of the RSA was done with both applications, and the resulting structures were compared against ground truth.

Participants evaluated each application with the System Usability Scale (SUS) [34], the User Experience Questionnaire (UEQ) [38] as well as the NASA Task Load Index Short (TLXs) [39]. These are description in Sec. 2.4.

#### 2.2.2 Procedure

Participants gave their informed consent. Participants were divided into four groups by ID. The first distinction was made on whether a participant received data with water-like noise. Furthermore, it is varies which application a user tested first, resulting in four conditions. The conditions were order of application and water noise.

The study procedure consisted of five steps. In the first step, participants would quantify their own previous experience and calibration measurements were made to setup the HMD. Afterwards, participants would be introduced to the first application (Desktop or VR) and after this initial training phase, the study data set would be loaded and the participant performed the task without help. Participants then evaluated the application using questionnaires. Lastly, these two steps would be repeated with the other application. For the full description of all steps involved in the individual phases, see App. B.

#### 2.2.3 Apparatus

In our experiment, we ran VRoot on an HTC Vive Pro HMD. In our tests, the application framerate was typically within 80 to 90 frames per second. The entire study was conducted in the “Virtual Reality Laboratory” in the Institute of Bio- and Geosciences 3 of the Forschungszentrum Jülich GmbH. The study was conducted sitting at the laboratory desk, facing the monitor. The VR software was used sitting by all participants. For considerations on whether to support or design a system for standing or seated setups, we refer to current literature [40].

The questionnaires as well as the NMRooting application were used on a desktop PC with a desktop resolution of 1920 *×* 1080 with mouse and keyboard. Our tests were performed on a PC with an Intel i7-8700K CPU, 32 GB of RAM and an NVIDIA 2060 RTX SUPER GPU.

#### 2.2.4 Participants

The user study data set was acquired from over 20 participants working on-site at Forschungszentrum Jülich. We have contacted potential participants, pre-emptively excluding anyone with either previous VRoot experience or knowledge of the goal of the study. Furthermore, we required normal or corrected-to-normal vision.

Total participation in the user study was *n* = 20. These include in total 15 male, 4 female, 1 non-binary and 0 other. Age distribution was almost uniform from 20 to 43 years, with a median age of 33. Self-reported experience using 3D applications was 4 participants with no experience, 8 users who reported using 3D applications at least once, 6 sporadic users and 2 experts. Self-reported experience using VR applications was 5: None, 5: Once, 7: Sporadic and 3: Expert. None of the participants had previous experience in the specific application NMRooting or the specific application VRoot.

### 2.3 Evaluation using FSPM Simulated Root Data

We are evaluating user-based extractions of root systems in the context of a virtual MRI scan. These virtual MRI scans were designed specifically for this study and the RSAs were simulated using the FSPM CPlantBox [5]. We calibrated the simulation for the task and slightly increased the inter-lateral distance for the first-order lateral roots. This RSA serves as a ground truth measurement. With noise that we typically see on a larger scale, such as Fig. 3.A, we modeled a smaller bean root system seen in Fig. 3.B and imposed a noise model on it.

**Fig. 3:**
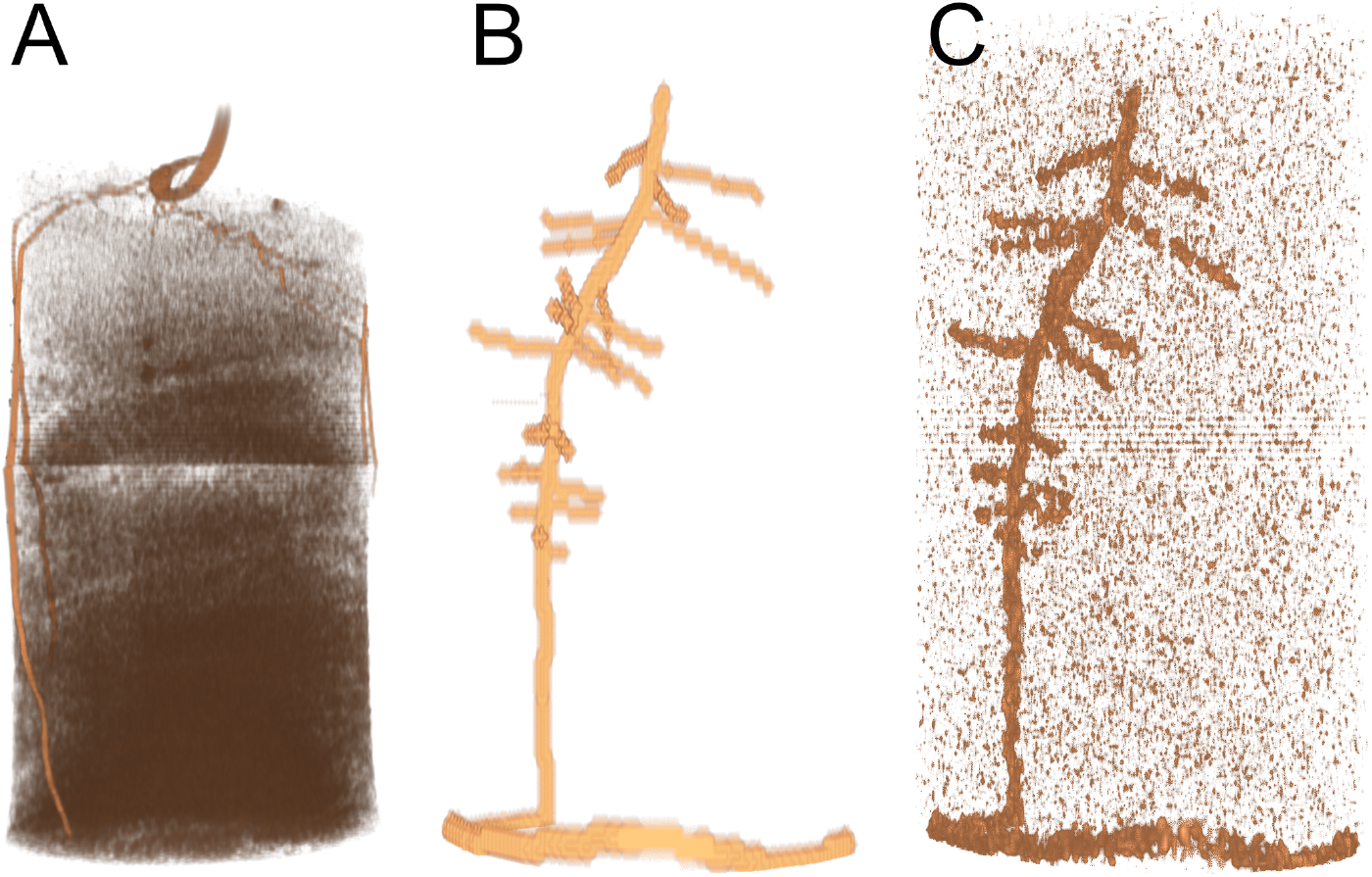
Side-by-side comparison of the study image data sets, scaled to image height as opposed to real height. A: We typically observe, depending on soil type as well as soil water content, highly ununiform noise. Locally, this might be expressed as smudges around the roots. B: Our simulation of a faba bean is being rendered with respect to the relative root length/diameter through a voxel. This image data set is the use-case for the noise-free participants. C: We added and subtracted noise features using a Weierstrass transformation. This results in slightly more complex, but uniformly complex, image data.

We computed the signal strength of the resulting MRI scan by using a simple heuristic based on the total volume of a root segment in a certain voxel. To avoid non-uniform task performance in the extraction task, the between-subject condition of water noise was chosen such that either the whole data set had noise effects or no part of the data. Within a soil cylinder of 1.5 cm diameter, a soil volume with water noise was seeded. The noise was locally scaled with the signal-to-noise ratio of 4.3. Noise addition/subtraction was followed by applying a Weierstrass transformation.

Since the original root system has been created by an FSPM, we can directly compare it with the manual extractions. This allows for an exact quantification of errors, which helps in assessing the factors in the decision-making process on what application to use for the pipeline. CPlantBox has been well-researched in terms of its stochastic properties [41], and is fit to be used as baseline for a user-based evaluation of extraction software. Additionally, any effects we would see in terms of the impact of the synthetic nature of the plant we would see in both applications equally.

As described in Horn et al. [11], to be able to confidently match between user-annotated RSAs and those generated by software or simulations, the correspondence between the architectures needs to be calculated. In this evaluation, we have used the distance-matching threshold of *d* = 15 in voxel units that was used by Horn et al [11]. Since any length of simulated root could correspond to one or more manually annotated segments, and likewise one manually annotated segment might correspond to more than one simulated root, we compute the full segment distance matrix as

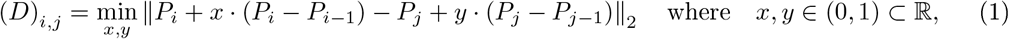

where *P*_*i*_ is the point coordinate of the *i*th segment, *x* and *y* are optimization parameters and the resulting distance matrix *D* only captures the minimal euclidean distance between two lines. This will result in a base distance metric that we use to match segments of different lengths. The matching process first assigns 1-to-n correspondences to simulated segments before it tries to match any unmatched manual segments to those simulated segments that were previously matched to exactly one segment. Organ-level label continuity is provided through matching roots. Two roots are considered matched if their cumulative distance *D ·* (*e*_*n*_ *× e*_*m*_) *↦ d*^(*m*,*n*)^ ∈ ℝ is the lowest among all other possible assignments with respect to the sets *e* of the segment indices.

We computed the accuracy based on root matching to ensure that the correct identification of roots is rewarded and to measure extraction differences that contribute to differences in root length. This topologically-aware accuracy is computed using a ground truth root set of *I*_*GT*_ and a set of traced roots *I*_*T*_, with a set of *I*_*c*_ := {*n ∈ I*_*GT*_ | argmin_*m∈*|*R*|_ *d*^(*m*,*n*)^ ∩ *d*^(*m*,*n*)^ ≤ *d*}, signifying the correctly matched roots.

To compute the measures for the correctness of the extracted RSAs, we first match the root systems to the ground truth. This ensures that there is topological information in the resulting scores. Root Lengths *L*_*n*_ generally refer to the total length of all segments corresponding to the root, namely 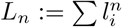 of the *i ∈ I*_*n*_ segments of organ index *n*. We use the ground truth *L*_*GT*_ as the reference for the scores, where *L*_*GC*_ is the total length of correctly matched roots computed from *I*_*GC*_ *⊆ I*_GT_. The root(s) that are matched to a root in the ground truth are a subset of the root set of the tracing, *I*_*T C*_ *⊆ I*_*T*_. It follows that the total length of false negative roots that were not traced is 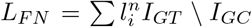. We especially highlight that the total length of correctly, which means matched, ground truth roots is not necessarily the same length as the sum of matched roots in the tracing, meaning that *L*_*GC*_*≠ L*_*T C*_ because *I*_*GC*_ *⊆ I*_*G*_ whereas *I*_*T C*_ *⊆ I*_*T*_. *L*_*F P*_ is the total length of false positive roots, i.e., computed from segments that were not present in the ground truth but present in the tracing. The recall value *R* is a measure that encapsulates how much (in length) of the root system was traced, defined as

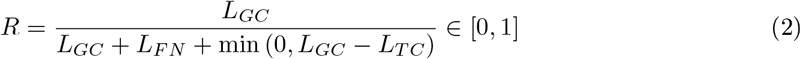

where we further penalize the tracing in cases where the extracted root length is smaller than the length of the ground truth. On the other hand, the precision value *P* encapsulates whether the manual extraction contains only as much length as the ground truth root system:

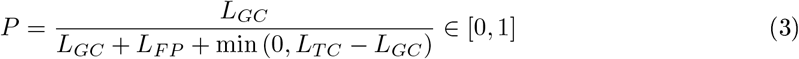

For the precision, we further penalize roots that were extracted correctly, but are too long in comparison to the ground truth. The precision and recall values are asymmetrical, decreasing with different metrics, but can be summarized by the symmetrical *F*_1_ score, which decreases with both false positives and false negatives:

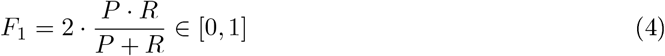

The *F*_1_ score is a comparison score that yields relatively similar values for different deviations from the ground truth. This score allows the comparison of applications that exhibit different characteristics, but due to its symmetrical nature, allows the comparison between them.

### 2.4 Measures for Application Comparison

The measures on usability of the software as well as user experience are difficult to measure objectively, and thus, questionnaires are typically utilized. These questionnaires include the System Usability Scale Questionnaire (SUS) [34], the User Experience Scale (UEQ) [38], and the NASA Task Load Index Short (TLXs) [39]. Due to the human interaction component of the system, we chose to use standard methods of evaluation with the added component of knowing the ground truth of a virtually generated MRI scan. Thus, we obtain a combination of subjective and objective measurements for assessing the software. We attached the full participant survey in the supplemental material.

The SUS is primarily aimed at quantifying the subjective user assessment of whether the given system is fit to help the user solve the problem. The resulting score is scaled within [0, 100] *⊂* ℕ. Commonly, the software is considered to rate well on this scale if it is above 85 [42].

UEQ scores are a way of evaluating the quality of the subjective user experience regarding basic descriptions of the software. The UEQ is a mix of adjectives that are presented in a contrasting manner. The adjective sometimes has overlapping meanings, and the general assessment of the software regarding these properties is very subjective. The NASA Task Load Index Short is a questionnaire to assess the subjective difficulty and strain on the user when completing the tasks. Users evaluated the applications in an online questionnaire, which we have attached to the supplemental material. We measured the extracted RSA, as well as camera data from participants, in addition to participants completing the questionnaire.

### 2.5 Data Analysis

We aggregated the data into two groups, based on the water noise condition. We computed the subjective scores per participant according to the respective guidelines. This applied to SUS [34], UEQ [38], and the TLXs [39]. We performed the data analysis entirely in Python. In tables or figures, we will refer to VRoot simply as VR, and to NMRooting as Water/No Water Conditions are referred to as +W or -W respectively. As such, VR+W refers to all data points of the VR software that have the water noise condition. In the following, total (T) refers to all data points. For hypotheses testing, our confidence cutoff is *p* = .05.

We tested all measures for normal distribution using the Shapiro-Wilk test, to ensure subsequent tests are informative and valid. For the sake of uniformity and comparability, we are using non-parametric tests in cases where not every condition is normally distributed. We chose the Mann-Whitney (Summed Rank) test for differences in median in cases where we do not find a normal distribution.

Statistical reporting includes the test statistic *t*(DoF), the critical value *p*, the effect size value (Cohen’s d [43], defined as 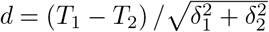 for differences in statistics *T*_*i*_), as well as degrees of freedom (DoF). In cases where ground truth is available, we test for a specific means using t-tests. The test values for normal distribution can be found in App. A. We use t-tests for SUS, UEQ, and TLXs. Mann-whitney tests will be used for total and average root length, *F*_1_ score, and inter-lateral distance.

## 3 Results

We omitted one subject (female, +W) from the study after the subject succeeded the task trace the taproot within the training phase but did not succeed with that task in the study phease. This means that we have 10 data points for the condition -W and 9 for the condition +W. The tracing and details on reasons of omission can be found in App. C.

### 3.1 Descriptive Data

We include box-plot descriptions of the relevant measures. Figure 4 shows the SUS scores over all conditions as well as the TLXs scoring and UEQ pragmatic quality. The SUS scores are scaled within [0, 100], though must not be understood as percentages. We have computed the median in instances of data points that have no normal distribution, which is indicated by the orange line in the boxplots. Data sets that contain a ground truth (simulation) value indicate this value with a red line. Data points outside of the inter-quartile range (1.5 *·* (*Q*_3_ − *Q*_1_)) have been included and marked as x-symbol.

**Fig. 4:**
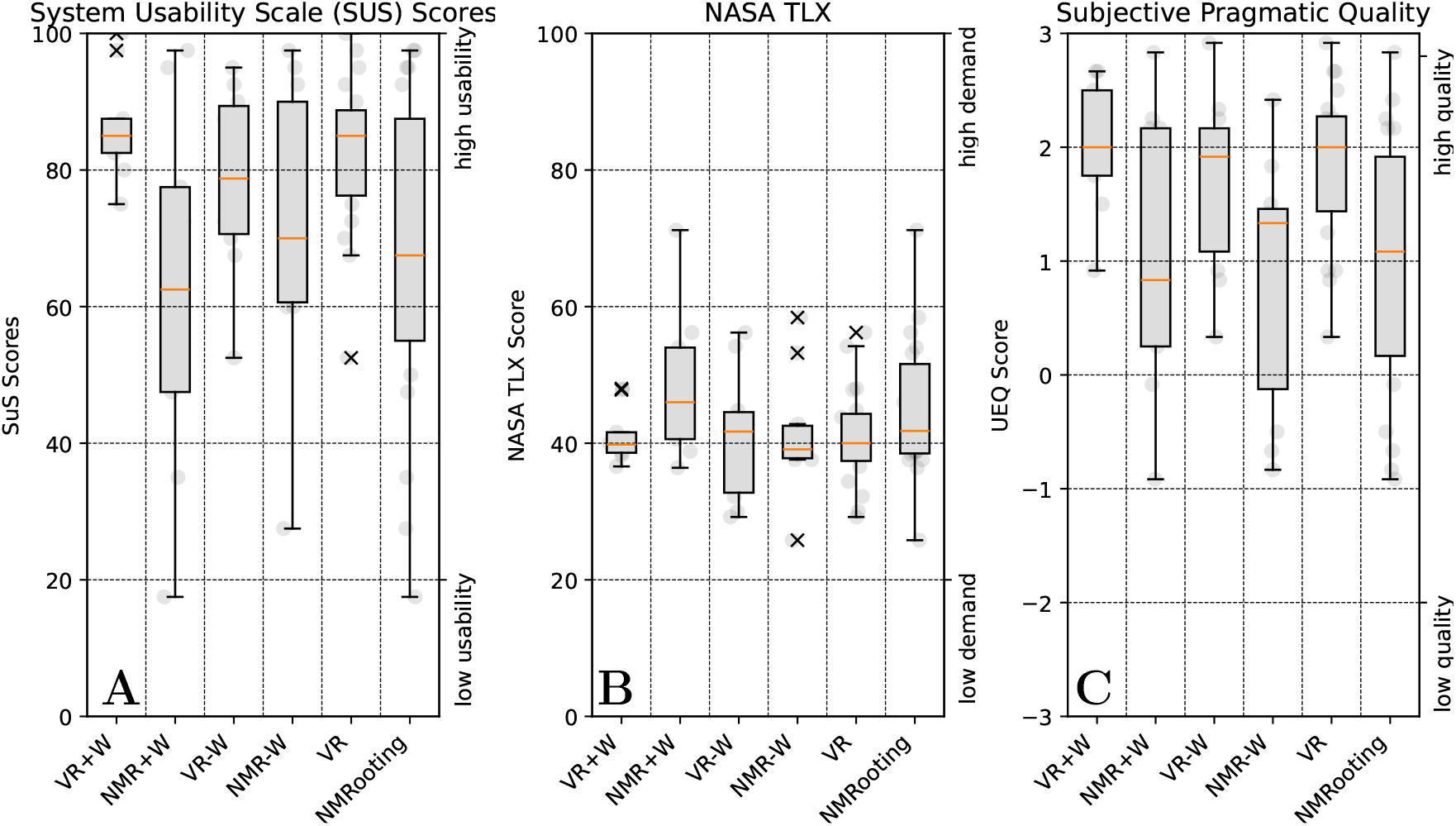
Box-plot with conditions on x axis and score on y axis, orange line is the median. A: Overview of System Usability Scoring across conditions. Scoring across order conditions (within-subject) was summed. B: Overview of Task Load Index Short Questionnaire score, sum of all questions except perceived success, which was inverted. C: Overview of UEQ participant scoring for the pragmatic quality of the data.

The within-subject condition of the order of the application was combined it served as balancing of the applications against learning effects. The subjective scores that are relevant to the applications can be seen in Fig. 4. Herein, we present the SUS, the Task Load as well as the pragmatic quality. Fig. 5 shows the accuracy scores of the data sets. For a more in-depth understanding of the individual effects, we further present relevant RSA measures in Fig. 6. Herein, we have a ground truth measurement for all conditions. Ground truth measures were extracted from the virtual MRI algorithmically, meaning that we extracted measures with the MRI mapping and voxelization in mind. For the comparison, we show as a red line the ground truth value from the actual simulated data set as opposed to the parameterization. For the number of lateral roots, we filtered roots of a length *l*_*i*_ *<* 3 [cm] to allow for the evaluation of the extraction without unnecessarily including tracing artifacts in NMRooting (seen in Fig. 7). The inter-lateral distance was calculated on the taproot only.

**Fig. 5:**
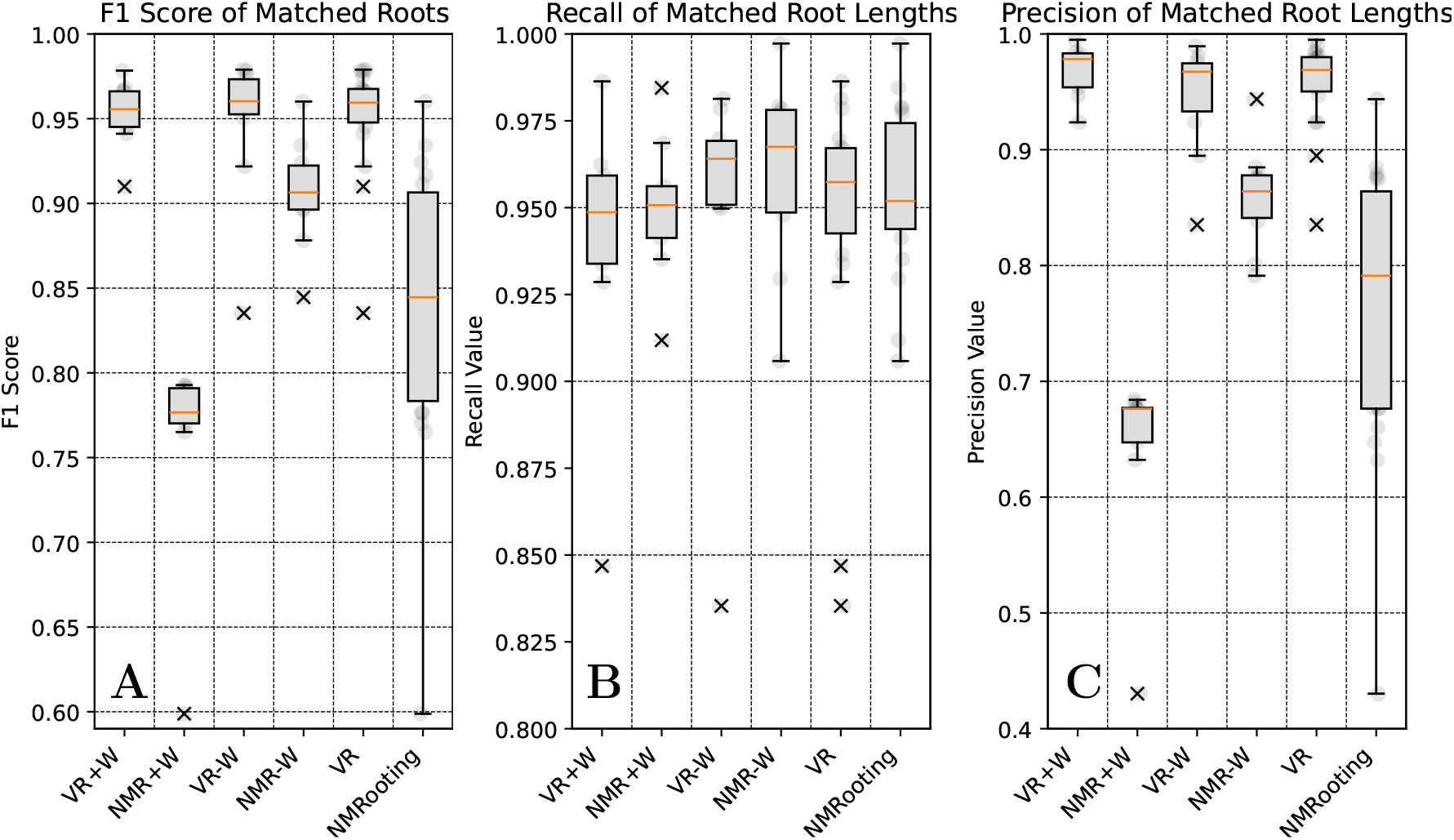
Box-plots showing the median (orange) and distribution of the data with outliers marked as x. A: *F*_1_ scores of the extracted root systems. B: Recall value of the matched root systems. C: Precision score of the matched root system.

**Fig. 6:**
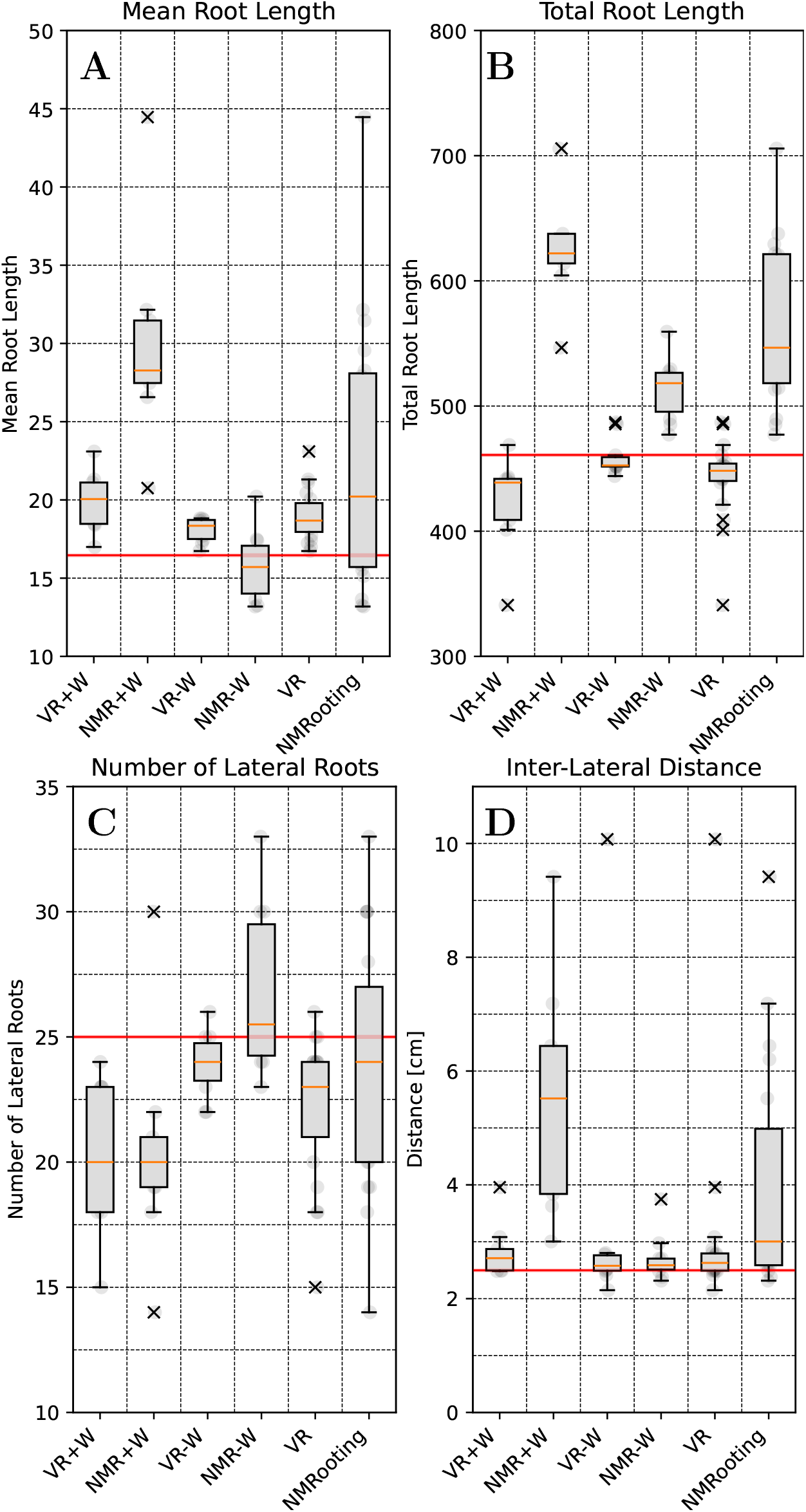
Box-plot graphs show ground truth in red, median in orange, and outliers as x. A: Boxplots of average root length B: Boxplots of total root length ∑_*i*_ *l*_*i*_ C: Number of lateral roots (*l*_*i*_ *≤* 4cm) D: Inter-lateral distance *d*_*i*_

**Fig. 7:**
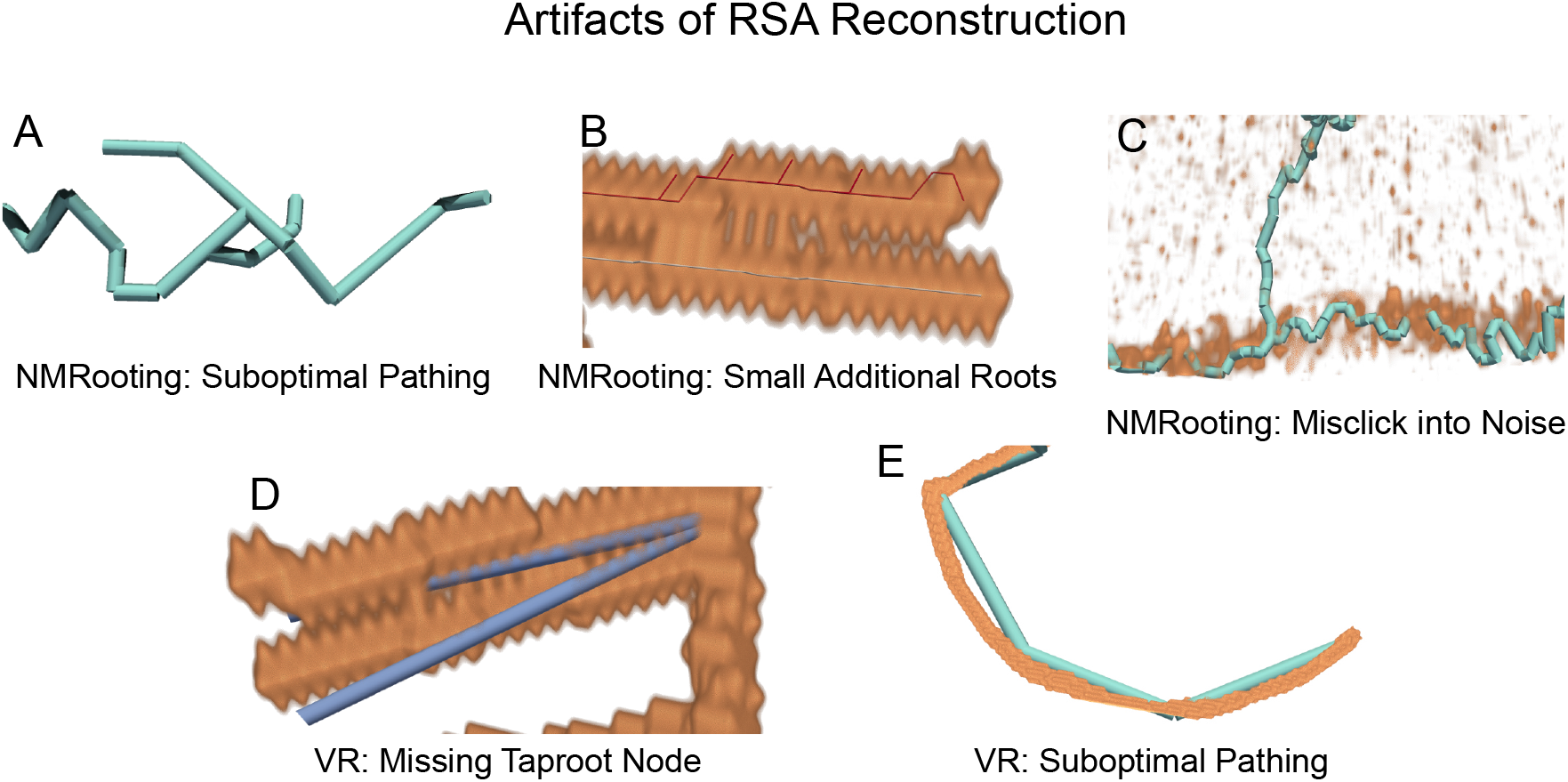
Artifacts of RSA reconstruction in each application that occurred with participants.

### 3.2 Results Regarding User Study Hypotheses

We summarized the statistics and hypothesis data in Tab. 1, which includes the most important data points. This section will give a brief overview of what results we measured on the hypotheses we had described in Sec. 2.2, and further indicators regarding ground truth comparison.

**Tab. 1:**
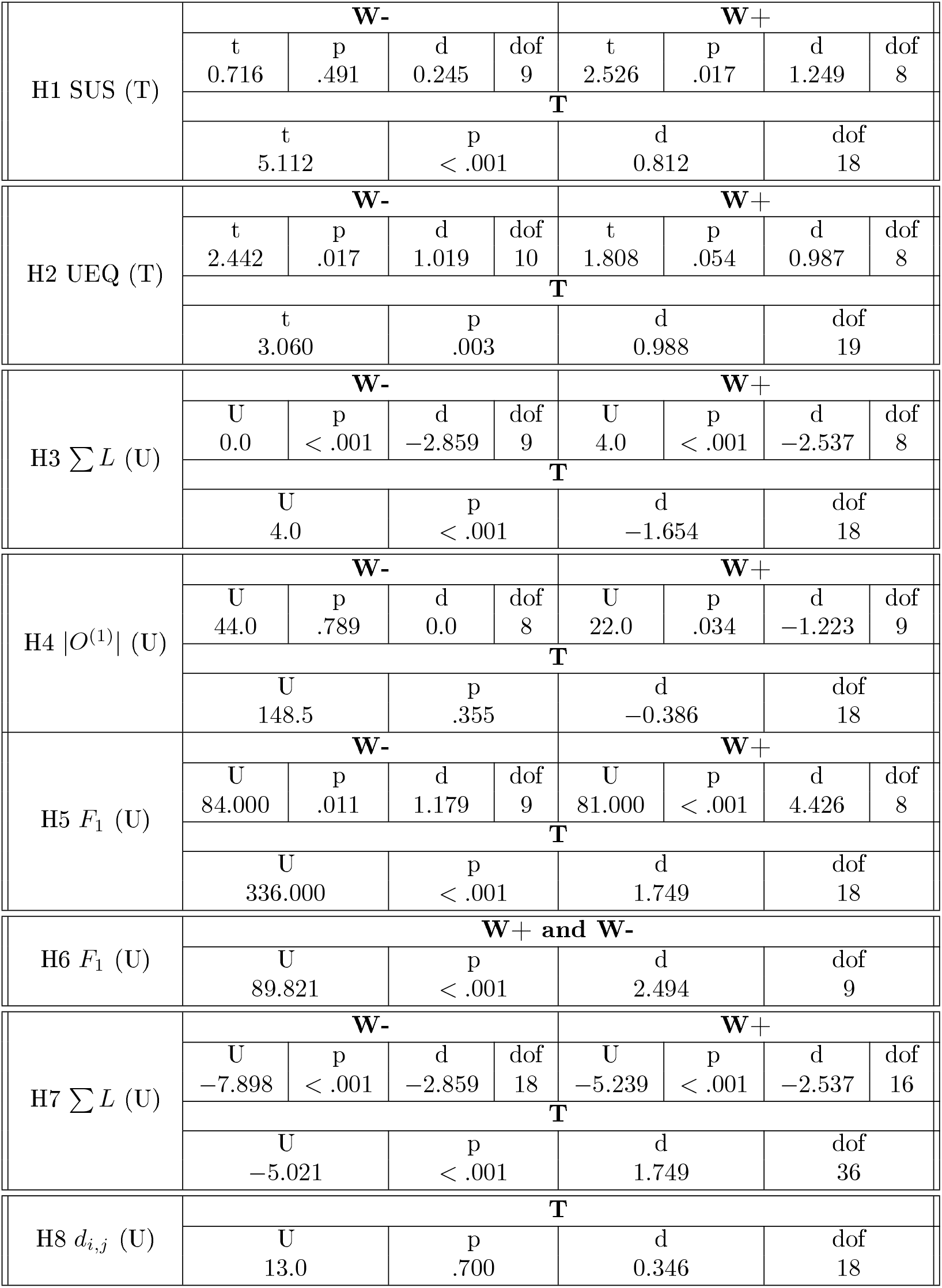
Hypotheses tests, reported with statistic *T* /*U*, critical value *p*, effect size *d* and degree of freedom (dof). We indicated what statistical tests were used by their abbreviation, namely U for Mann-Whitney U-Statistical Test and T for T-Test.

#### SUS

We observe an average usability of 82 for VR and 67 for NMRooting. VRoot has scores significantly better than NMRooting as measured using a one-sided t-test (H1). Notably, VR+W has a average usability of 86 and NMR+W of 62, with tests showing significant improvement by using VR, which is not the case for the comparison VR-W *>* NMR-W.

#### UEQ

Pragmatic experience that can be extracted from the UEQ is the average of the three statistics Perspicuity, Efficiency, Dependability. VR scored 1.841 on average and NMRooting scored 0.912, resulting in a significant difference (H2).

#### Root Length

The length measures of the root system are average and total root length, seen in Fig.6.A and B respectively. We computed the differences in extraction quality regarding these two measures, allowing for an assessment of overall extracted biomass and the correct identification of individual roots. The average and total root length contain useful information about what challenges participants encountered during the annotation. We observe a difference between NMRooting and VR in terms of extracted root lengths (H3).

Deviation from ground truth was tested as well as the difference between the methods. The extraction of the average root length was, on average, correctly done in NMR-W, by a single-sample t-test (*t*(9) = −0.979, *p* = .353). The total root length extraction in VR-W yielded no significant difference to ground truth, yielding *t*(9) = −0.426, *p* = .680. Other conditions, including aggregates, yield significant differences. Differences between the applications exist in the case of the individual conditions (+W/-W) as well as the aggregate. We furthermore observe that the extraction of the total root length yields larger differences in the +W condition.

#### Root Topology

Computing the extracted number of laterals that have a minimum length of 3 cm, we find that only NMR-W found the correct number of lateral roots. For the branching density (H4), we observe a difference between the extracted number of lateral roots in the +W case, but not in the -W case nor in the aggregate case.

*F*_1_: We find that the *F*_1_ score of the extracted root systems is significantly higher for VR in both +W and -W conditions (H5). Additionally, we find an increase in the difference between the applications once water noise is present (H6).

#### Influence of Water Noise

We observe a significant increase in the difference between the applications when water noise is present. This is the case for the average root length, the total root length, and the *F*_1_ score. The computation for the *F*_1_ score test required random pairing between individual scores.

#### Inter-Lateral Distance

This measure is defined as the average distance between two consecutive lateral organs *O*_*i*_, *O*_*j*_. VRoot users correctly extracted the inter-lateral distance, tested using a two-sided Mann-Whitney-U Test. NMRooting yielded a significant difference to ground truth in the +W condition, by single-sample t-test with *t*(8) = 1.619, *p* = .002 and an effect size of *d* = 1.575. There were no significant differences in both VR cases as well as the NMR-W case.

## 4. Discussion

We postulated that the VR software would yield a different usability, which has been confirmed in H1 for W+ and overall use of the software. The effect was much smaller for W-, resulting in a very small effect size and no significant difference. However, while no significant difference was measured, there is furthermore no indication that classical applications perform better in any of the study conditions. The overall measured variance for the measurement of pragmatic quality (H2) was fairly high, see Fig.4.B, resulting in no significant difference between the applications in the W+ condition.

Objective measurements were more uniformly successful, with the notable exception being the detection of the correct number of lateral roots. In that metric, all median extractions were below ground truth, with the exception of NMR-W, which was slightly higher. Interestingly, NMR+W extractions consist of less roots overall, which might be a result of the noisy data inhibiting the participant’s ability to extract roots. We confirm other findings, such as Selzner et al. [12] and Horn et al. [11] about the impact of noise on the RSA reconstruction.

### 4.1 Task Execution

Generally, task execution posed no problems for users. Users needed a few minutes to get accustomed to using the HMDs. Though pre-emptive measurement and calibration were done for the inter-pupillar distance, some users reported issues with depth perception, or depth-perception issues became apparent during the training phase. Though explicit introduction and prompting to repeat a certain interaction were done during the training phase, some users did not make use of all available options, particularly navigation, during task completion. This occurred in equal parts with NMRooting and VRoot.

### 4.2 Interpretation of Results

The simulated root contained certain artifacts that would have made it fairly hard to trace for new users. Particularly, we note that the matching-based *F*_1_ score is remarkably good for the VR software, which is in part due to users matching the correct root length for the individual organs successfully. The spatial distribution of the root system is more obvious in VR, which reflects in the root systems that were drawn, even under the condition that users do not edit root nodes that were already placed. The restricted set of functionality was mitigated by the introduction of a training session, during which the procedure was explained, leading to most participants already pre-planning the taproot in a way that made it possible to achieve a high accuracy.

One aspect of the results we would like to highlight is the fact that with no water noise, the VR application still yielded better results in terms of matched length-based *F*_1_ scoring, see Fig.5.A. Furthermore, the differences between the individuals were not very high, resulting in a standard deviation for the VR conditions of 0.02 (W+), 0.04 (W-), and 0.03 (T) respectively. However, thi is partially due to the fairly ‘destructive’ nature of the *F*_1_ score, leading to different user-based tracing errors to result in a similar score. The recall value *R*, as seen in Fig. 5, is fairly uniform across applications, with slightly more “undertracing” done in the VR application. The *F*_1_ score and particularly the precision *P* was low (but not extremely so) for NMRooting in the +W condition. The primary reason for this was the fact that each time a user clicked into the data and did not continue a root from the tip but rather from the closest segment regarding the signal strength, this induced a small lateral root at that point that users might have missed. For the actual comparison of the amount of lateral roots, we discarded shorter roots to avoid comparing against this technicality. We note that the *F*_1_ scores closely follow the power law distribution, tested by means of Cressie-Read test for goodness-of-fit at *χ* = 0.021 and *p* = 1.0.

There is a significantly higher spread in the perceived system usability measured from NMRooting in the +W condition as well as overall. Tested variances were significantly higher according to the F-test for the overall condition (*F* (18) = 4.330, *p* = .002) as well as +W (*F* (8) = 11.493, *p* = .001) but not for the conditions -W (*F* (9) = 2.571, *p* = .087), even though this condition does fail the test for equal variance (*F* (9) = 2.571, *p* = .175). Between the VR+W and VR-W conditions, there is an increase in variance that is, while not significant, at least notable. Moreover, the inter-lateral distance has a large variance in the NMR+W condition due to several artifacts. We will note that, for the estimation of the inter-lateral distance, while there were no significant differences in VR+W, VR-W, and NMR-W, that most users (68% for VR and 84% for NMRooting) over-estimated the inter-lateral distance compared to the ground truth. We cannot make assumptions on the applicability of this regarding a real MRI scan, but our findings provide some insight into user bias when extracting parameters, particularly for FSPM simulation, from 3D imaging data. The average relative error for inter-lateral distance for VR users was 0.234 and for NMRooting it was 0.617.

The average and total root length is, in part, influenced by the presence of water noise, in both applications. NMR+W exhibited a larger total root length while still yielding a lower than ground truth number of laterals. However, the increased inter-lateral distance partially relates to misidentification later in the data. In VR, the presence of water noise caused under-identification of roots. We will note, that in the simulation data, there was one very thin root that would have been very hard to identify. There was a systematic issue with users not being able to identify that root and thus, the VR-W case was lower in median regarding the number of lateral roots users were able to identify.

### 4.3 Artifacts of the RSA Reconstruction

Our general results show an improvement in the extraction quality using VR. More specific phenomena can often be explained taking into account observations from the study or by closer inspection of the data. There are a few instances of false positives within the NMRooting annotation that can be attributed to users clicking on surfaces they did not intend to. It is important to note that during the eventual study task, no further assistance was provided unless prompted, as opposed to the training task, which included guidance and repetition until no mistake was made.

This effect was exasperated by the presence of water noise. Some users corrected their annotation to a certain degree, resulting in fewer false positives but still suboptimal pathing, as seen in Fig. 7.A. In contrast, a case of supoptimal pathing in VR, which directly results from a coarser user interaction, is seen in Fig. 7.E. A total of 5 users attempted to separate the proximal roots, which yielded a few perfect annotations, but also artifacts such as Fig. 7.B, which includes a few small roots that are due to participants progressing the lateral by clicking in smaller steps, such that the algorithm does not use the connecting signal between the roots to path to the point indicated by the user. If water noise was present, a few users misclicked into the water volume during the study task but failed to remove such a tracing at a later point, as seen in Fig. 7.C.

While the VR application had better *F*_1_ scores on average, ultimately more training is required for users to produce high-quality RSA reconstructions. Some participants struggled with depth perception in VR, which occurred in approximately equal parts in people with corrected vision and normal vision. This is likely an experience effect that would only be resolved by further use of the VR system - participants with previous VR experience did not encounter this. Targeting objects in the virtual scene as well as the correct placement of segments was challenging for new users.

The generally low variance of individual scores might be an indicator of a systematic effect that is uniform among individuals. In the case of VR, the inability to edit nodes was likely a large contributor to the overall score of participants. This caused issues in cases where there was a node missing in the taproot, as exemplified in Fig. 7.D. In the case of NMRooting, participants were told to manually trace the nodes and were able to delete nodes that were mistakingly inserted. The low score in NMRooting is mostly due to smaller effects accumulating to a lower score in total, including supoptimal pathing, misidentification of roots, as well as the presence of proximal roots and water noise in certain conditions.

### 4.4 Explanations for Length Differences

VR users tended to estimate the ground truth total root length correctly when dealing with noise-free data. However, there is a slight indication of overestimation of the individual root length within noisy data, as found in Fig. 6. VR users might not have drawn the taproot correctly, resulting in a slight overestimation of average root lengths, but still an underestimation of the total. It has to be noted that the task was performed sitting, which meant that users were forced to use the navigation in VRoot as an alternative to bending down. Some users chose to trace the root system less effectively as parts of it were out of reach, resulting in a higher variance of the total length measurements. A few users requested to be able to stand, but for the sake of uniformity, we did not allow this.

On the other hand, the large overestimation in average root length in NMRooting was in part due to supoptimal pathing, while the overestimation in total root length was caused by false-positive lateral roots, which is indicated by the shift in distribution in the inter-lateral distance, as seen in Fig. 6. We further investigate this by filtering the roots for only true positive identifications and subsequently compute their length difference, as shown in Fig. 8. While we do observe a higher-than-zero length extraction, by means of double-sided t-test with *t*(198) = 2.379, *p* = .018, *d* = 0.169, we observe a significantly higher variance in the +W condition of the desktop software (*F* (192) = 0.651 and *p* = .001). We believe that a combination of issues, most notably inability to effectively navigate in a desktop setting, caused the differences in the +W condition.

**Fig. 8:**
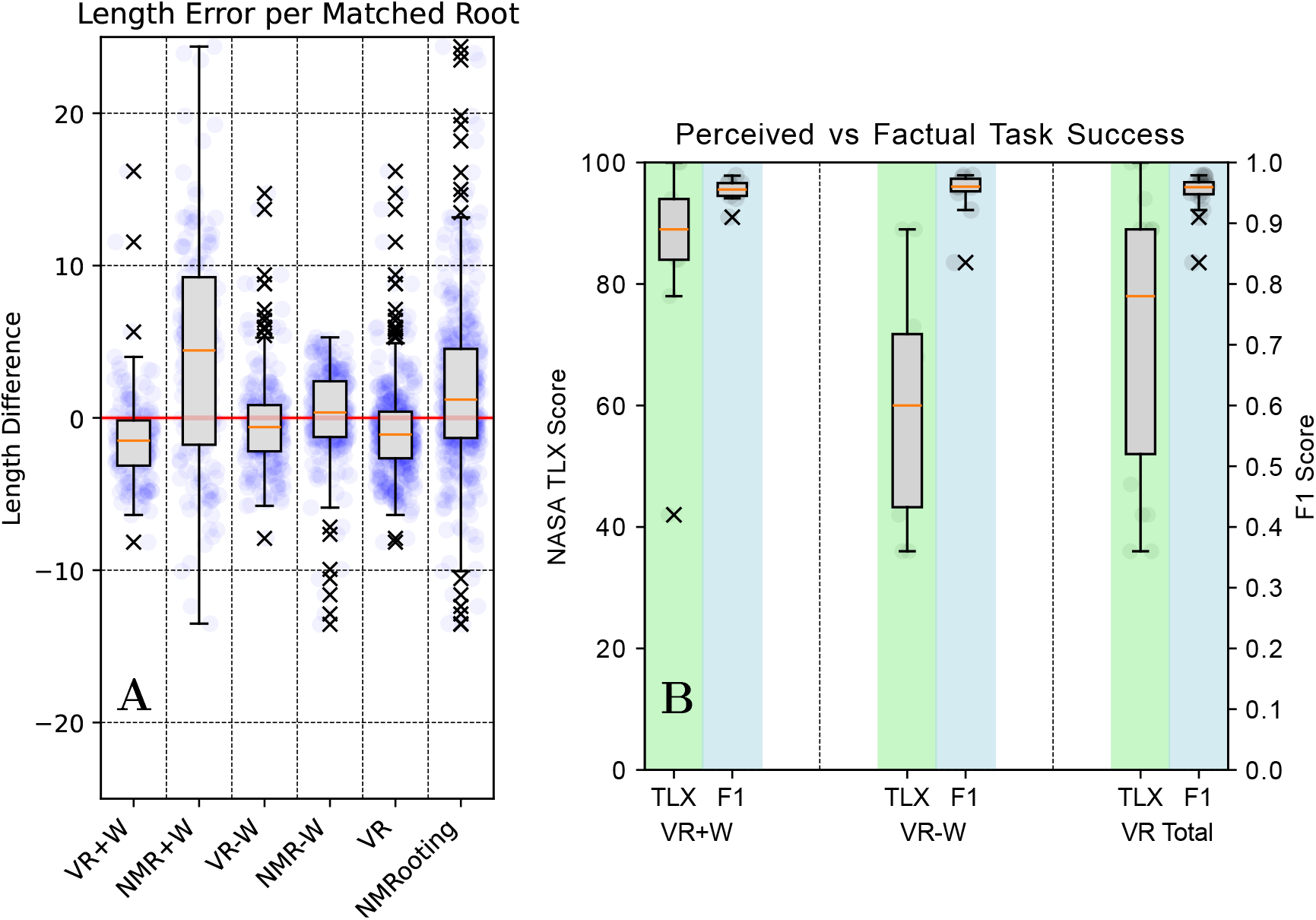
A: Box-plots of length differences between ground truth and correctly annotated roots. Ground truth is red, median is orange, raw data is indicated blue, outliers are marked as x. B: Comparison of self assessment of extraction quality in VR to actual extraction quality.

### 4.5 Subjective Measures

The quantification of responses to the questionnaires yielded mixed results. Particularly, we want to highlight the increased usability through the use of VRoot. While there was no significant increase in the no-water condition, as seen in Fig. 4, there was a larger spread of responses for the NMRooting software, resulting in a higher average usability score of using VR in comparison. NMRooting generally had a larger spread in responses in contrast to the more unified responses for VR both in total (*F* (18) = 4.330 and *p* = .002) and in the +W condition (*F* (8) = 11.493 and *p* = .002). The TLXs yielded no significant difference between NMRooting and VRoot, but it did showcase a higher variance in the -W cases.

The user experience is challenging to compare, as certain measures (such as novelty) are not appropriate in the assessment of the desktop software. This is especially true since participants will already have the expectation of using VR software even if using the desktop software first, leading to influences measurements of those metrics.

The task, while the root was generally simple with only a taproot and 25 laterals, there were particularities about the data that caused issues for certain participants, especially when using NMRooting. However, artifacts like proximal roots, or smaller roots further down the taproot, have gone unnoticed to some participants. We will note that there was no difference in the individual ratings depending on the order of applications tested. Generally, it is quite natural that the tasks would appear equally demanding between the individual conditions. Interestingly, the VR+W condition has the smallest variance, indicating a more universal agreement even between the within-subject conditions.

One aspect of the TLXs questionnaire is the question on the self-assessment on whether the task was completed. This assessment is shown in Fig. 8.B. The *F*_1_ score shows that users performed similarly whether there was water noise present, or not. However, self-assessment of the accuracy was much lower than the actual accuracy with no water noise present. This is likely less a self-assessment and more a comparative assessment depending on how easy the problem appears to be solvable with automatic means. We highlight this to underline the issue that the subjective assessment of data extraction done by users is seldomly representative of the actual data quality. While our users estimated their own performance as worse than it actually was, this is a more general issue, as manually annotated public data sets also contain label errors, which generally are assumed to be perfectly labeled [44]. However, manual annotation work is incredibly valuable, especially in plant science. The human self-assessment of data quality measured through manual means, especially if used as training basis, is seldomly accurate.

### 4.6 Combination of Methods

Even if a virtual reality workflow improves the quality of extraction, manual tasks remain more tedious and time-consuming than automatic extraction. Our workflow is ultimately aimed at correcting rather than tracing.

VRoot functions best with a mixture of high throughput pre-tracing and features that assist with the analysis of root architectures as the basis. In the future, we would like to combine automated, semi-automated, and manual tracing methods. In the case of NMRooting, this would require the introduction of additional interaction metaphors suitable for tasks that are voxel-based. There are other methods, such as TopoRoot [10], that could improve the manual extraction pipeline. This would be in line with published literature on similar topics, namely by Zeng et al. [29], who improved the VR application developed in Usher et al. [14] through the use of topological features.

## 5. Conclusion

In this work, we presented a pipeline to extract RSAs from MRI images using VR. We established the need for a more immersive manual analysis tool for complex data sets and showed the advantages of using VR to enable new users to achieve high-quality reconstructions faster. We evaluated the use of our VR software in comparison to contemporary desktop applications, and semi-automated analysis. Furthermore, we quantified how well participants with a uniform knowledge base performed in these tasks, both on the desktop as well as in VR. Our results show an increased usability and accuracy through the use of VR for manual root workflows, especially in instances where automatic tools need more assistance. This enables the analysis of root systems in more diverse soil conditions. Here, immersive annotation is a very valuable method in 3D root image analysis, and helps to increase the variety of analyzed data to more soil types and soil water contents. In the future the goal is to combine the tools offered by our VR application with the benefits of an automatic extraction to provide a user-friendly and fast workflow to correct automatic tracing results.

## Supporting information

Survey as PDF and XML

## 6 Ethical Statement

The study conception and planning got approval from the ethics committee of Trier University. Participants provided their explicit consent and were instructed on how to contact us to have their data deleted should they wish to do so. Data is stored and published anonymously.

### 6.1 Author Contributions

This work was primarily authored by Baker. All authors discussed the results, provided feedback, and contributed to the text of this work. Study conception and execution was done by Baker and Zielasko. The implementation of the VR application and development of its features was done by Baker, Selzner, Göbbert, Zielasko, and Schnepf. Scientific counsel and directions for data analysis, study procedure, as well as results was provided by Selzner, Scharr, Riedel, Hvannberg, Schnepf, and Zielasko. Funding for hardware was provided by Schnepf and Göbbert. Zielasko has primarily supervised this project and guided implementation of its tasks.

### 6.4 Data Availability

The data needed to reproduce the results of this work has been uploaded to 10.26165/JUELICH-DATA/B9SB0S. We have included a meta-description of the approach but encourage researchers aiming to reproduce our results to reach out. The Software VRoot will be found on Github: dhelmrich/VRoot. The questionnaires have been attached to the supplemental material. We have published a video description of the software, seen in 10.6084/m9.figshare.26003494.

## A. Normal Distribution Tests

The shapiro-wilk test for normal distribution tests whether the data is drawn from a normal distribution. This is a test where the alternative hypothesis is exhibiting a normal distribution, meaning that the *H*_0_ hypothesis is that *X* ≁ *N* for some normal distribution *N*, referring to a significant difference to data produced by a normal distribution. For comparative measures, the residual of the two statistics, *R* := *T*_1_ − *T*_2_, needs to be normally distributed, meaning that the difference between applications is tested as opposed to the statistics *T* of each application itself.

As seen in Tab. 4, there are some instances of data where a normal distribution is not present, which are the +W conditions for the root lengths, resulting in a difference from a normal distribution for the total data. The inter-lateral distance *d*_*i*_ does not exhibit normal distribution for the -W condition.

**Tab. 2:** Goodness-of-fit tests for normal distribution on different measures, including combined measures. The pairing of residuals is always within subject.

| Measure | Water |  | No Water |  | Total |  |
| --- | --- | --- | --- | --- | --- | --- |
|  | t | p | t | p | t | p |
| SUS | 0.902 | .264 | 0.959 | .783 | 0.944 | .311 |
| UEQ | 0.939 | .577 | 0.938 | .497 | 0.897 | .051 |
| TLXs | 0.933 | .515 | 0.953 | .713 | 0.976 | .897 |
| TRL | 0.798 | .019 | 0.870 | .101 | 0.744 | < .001 |
| ARL | 0.798 | .019 | 0.870 | .101 | 0.744 | < .001 |
| $F_1$ | 0.722 | .002 | 0.931 | .462 | 0.909 | .072 |
| $d_i$ | 0.915 | .357 | 0.490 | < .001 | 0.901 | .051 |

**Tab. 3:** Goodness-of-fit tests for individual statistics for the testing against ground truth measurements, for each VR condition.

| Measure | VR+W |  | VR-W |  | VR |  |
| --- | --- | --- | --- | --- | --- | --- |
|  | t | p | t | p | t | p |
| SUS | 0.902 | .264 | 0.959 | .783 | 0.944 | .311 |
| UEQ | 0.932 | .504 | 0.961 | .790 | 0.953 | .074 |
| TLXs | 0.877 | .157 | 0.907 | .259 | 0.913 | .074 |
| TLR | 0.679 | .001 | 0.777 | .011 | 0.719 | < .001 |
| ARL | 0.986 | .988 | 0.879 | .129 | 0.928 | .159 |
| $F_1$ | 0.908 | .308 | 0.686 | .001 | 0.720 | < .001 |
| $d_i$ | 0.756 | .006 | 0.443 | < .001 | 0.404 | < .001 |

**Tab. 4:** Goodness-of-fit tests for individual statistics for the testing against ground truth measurements, for each NMRooting condition.

| Measure | NMR+W |  | NMR-W |  | NMR |  |
| --- | --- | --- | --- | --- | --- | --- |
|  | t | p | t | p | t | p |
| SUS | 0.963 | .824 | 0.884 | .146 | 0.931 | .160 |
| UEQ | 0.920 | .394 | 0.918 | .302 | 0.944 | .289 |
| TLXs | 0.839 | .043 | 0.924 | .390 | 0.882 | .019 |
| TLR | 0.744 | .007 | 0.814 | .014 | 0.714 | < .001 |
| ARL | 0.930 | .453 | 0.854 | .083 | 0.885 | .026 |
| $F_1$ | 0.539 | .007 | 0.979 | < .001 | 0.890 | .032 |
| $d_i$ | 0.949 | .687 | 0.771 | .006 | 0.807 | .001 |

These effects were likely caused by a non-uniform error that is present within the data, as opposed to the subjective measurements that are mostly centered around an average value, the objective distributions are non-symmetric regarding the differences between the applications.

## B Study Procedure

In the questionnaire for the study, participants were informed about what data would be stored and how. Participants gave informed consent for the participation in the study. Participants were assigned an ID at the start of the study, and we measured their inter-pupillar distance for the purposes of callibrating the HMD to their respective measurements. Depending on the ID, participants then used either the application VRoot or NMRooting to extract a root system. For both applications, participants were verbally explained the functionality of the applications and what features they were to use to extract the root system. Each participant was tasked with performing all actions on a training data set that were required later on, which included navigation, tracing, and deletion in the case of NMRooting. After task completion with the study task, during which no intervention took place, participants filled out the questionnaires attached in the supplemental material. Participants will then repeat the process with the other application.

The study aimed to assess the direct interaction with the system in cases in which manual annotation was necessary. As such, participants were told to only use these tools to ensure comparability, as otherwise the task would be more complex, and user interaction would require more functionality and a deeper understanding of the application. In the VR application, participants were taught the draw root functionality as well as data set navigation techniques to ensure that all of the data set is reachable. Successful completion of the VR training task required the following steps:

1. Move head position in VR
2. Rotate the root system
3. Move the root system up/down
4. Select a node
5. Deselect a node
6. Draw a node
7. Deselect a node and draw a lateral

In NMRooting, we restricted the functionality to the tip-to-tree option, with the correct starting point already being set pre-emptively. Users did label the root topology manually. Successful completion of the NMRooting training phase required the following steps:

1. Turn the view
2. Zoom in/out
3. Pan the view (lateral movement respective to the screen view)
4. Click into the data to add a root
5. Click into the data to delete a root
6. Open the root order menu
7. Label roots by their root order (here just taproot and first order lateral)

Our previous tests yielded an average completion time for the tasks of 45 min. From a previous test with the application with mixed, i.e., expert and non-expert, participants (*n* = 16), we concluded that one hour would be sufficient for untrained participants. While we did not provide a timing nor a time for task completion in the individual conditions, we did allocate time slots for participants which were an hour long.

## C Outlier

The outlier data set, exemplified by the VR performance of the participant. While the participant was able to annotate both the taproot as well as annotate lateral roots in the training task, the participant has not retained the knowledge of the annotation steps and produced an annotation (red) as shown in Fig. 9. It is unclear whether this was caused by the presence of water noise.

**Fig. 9:**
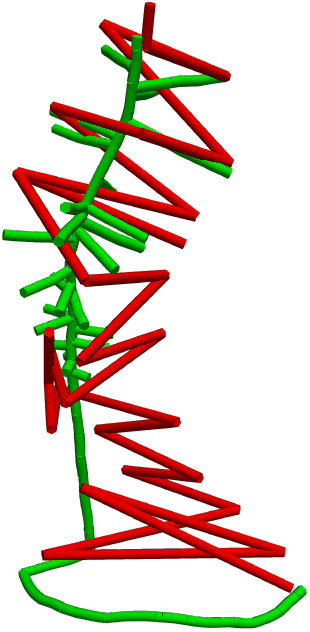
Removed outlier data set after subject failed to trace the taproot. While the cause of this issue is unknown, the participant did trace a fully functional root system in the training stage of the task, which included being prompted to place nodes on the taproot of the root system.

