## Supplementary material for "VRoot: An XR-Based Application for Manual Root System Architecture Reconstruction": Survey as PDF and XML: survey.pdf

### 3D Root Architecture Tracing Usability Study

There are 45 questions in this survey.

#### Study: Assessing the correctness and usability of Virtual Reality for Root Data Analysis

Welcome to the user study of this collaboration project and we thank you for your interest in furthering research. This study will take approximately 40 minutes

### Study Procedure

After you read this information sheet and you provided written consent to the collection of your data as well as your participation in this study, you will receive a random ID number we will use to anonymize your data. You will be the only person that knows this ID number. You will be asked to fill out a form about yourself and your previous experience in the relevant techniques
(root data analysis, plant image analysis, 3D applications, and virtual reality).

The study consists of two parts: In one part, you will be asked to trace root systems with the desktop analysis software NMRooting. You will have a few minutes to accustom yourself to the software and the means of interacting with the data. You will be shown a root system scan (like an MRI scan) and your
task is to trace the root system to the best of your abilities. The second part will include the use of the head-mounted display (HMD). This display system is a wearable Virtual Reality device with which you can see the scene and interact with it. You will have a few minutes to accustom yourself to the headset and the means of interacting with the scene and its data. You will be provided with an interaction sheet explaining the different buttons of the hand-held tracked devices. In this part of the study, you will be shown similar a root system with the task of tracing it. There will not be a functional difference between the two root systems that you will be tracing, they will be scans from the same batch of plants, but two different root systems. During the study, we will track your head and controller interaction with the VR device. We will also track your mouse movement as well as click interactions with the desktop setting.

The study as a whole will take approximately 40 minutes.

### Risks

This study does not pose a risk to your health and well-being. There is a chance of discomfort during the use of Virtual Reality devices, such as an HMD. If at any point in the study, you feel nauseous and want to stop the experiment, please inform the experiment staff. You can also ask to end the study at any time and will be helped by the staff. Furthermore, there are grid lines displayed in the virtual reality scene that marks the VR movement area. Please stay within those grid lines.

### Anonymity and ongoing consent

Participation in this study is entirely voluntary you can revoke your participation at any point in time without providing reasons. This will not have negative consequence

### Data Protection

Any information that is obtained during this study will be kept confidential. You are assigned an ID number known only to the researchers, with information disclosing your identity not to be released without your consent. Participants will not be identified by name in any reports of the completed study or any data that will be shared with other researchers. This survey data will be stored electronically on password-protected storage devices or secure servers and kept for at least 10 years. Data might be kept indefinitely. Aggregate data (but not consent forms or any other identifiable information) may be uploaded to an open-access data repository for others to use, and thus may be kept indefinitely; this data will be de-identified, meaning any direct and indirect identifying information will be removed or transformed. The results of this study may be reported in a graduate thesis, online on websites, and may also be published in other ways such as articles, books, and conferences. Additional research may be conducted where the data is used for future analysis within a different study, with any identifying information removed (de-identified) from the data set. The personal data that you are giving us to contact you will be treated confidentially and not be connected to your study data or personal Participant ID. Your personal data will be stored securely either physically locked away (if on paper) or on a password-protected server (if digital) within the Forschungszentrum Jülich GmbH.

### Contact Data

**Studienverantwortlicher**
**Fachbereich IV**
**Informatikwissenschaften**
Human-Computer Interaction
Dr. Daniel Zielasko

Behringstraße 21
54296 Trier

Tel.:"0651 201 3201


**Versuchsleiter**

Forschungszentrum Jülich GmbH
Jülich Supercomputing Centre
Dirk Norbert Helmrich

Wilhelm-Johnen-Straße
52425 Jülich

Tel.: +49 2461/61-1792


#### I have read the above information and agree to the terms of the study. *

Please choose **only one** of the following:

- ◯ Yes
- ◯ No

### Informed consent

#### Please read this section carefully.

By acknowledging this section, you agree to the following:

- You have been given written information about the topic and procedure of the study that you are about to take part in. Any questions you had have been answered to your satisfaction.
- You have been informed that you can withdraw from the study at any time without having to fear negative consequence

You consent to the storing and use of your data according to the agreed-upon conditions. The recording of your data will be done using a pseudonym which only you can connect to your person. This means that no one else can relate your study data to you during the evaluation process. You know that you can request the deletion of your data at any point without this leading to negative consequences for yourself. You agree to the use of your de-personalized data for research purposes and their storage for at least 10 years into the future. You furthermore agree to the secure storing of your personal contact data for the sole purpose of being contacted about the study that you am about to participate in.

*

❗ Choose one of the following answers
Please choose **only one** of the following:

- ◯ Yes, Proceed.
- ◯ Get me out of here.

### Demographic Information

Please enter your Code. You will have received a **code from the study supervisor**. Note that this code is important and you will be **asked for this code three times** in total. *

Please write your answer here:

Please state your age. *

Please write your answer here:

Please state your gender *

Choose one of the following answers

Please choose **only one** of the following:

- Female
- Male
- Non-Binary
- Other

Please rate your previous experience with the analysis of any plant related data. This can include organ counting, parameter extraction, root length measurements through root washing or rhizotron images, etc. *

Choose one of the following answers

Please choose **only one** of the following:

- No experience
- At least once
- Sporadic experience
- Expert

#### Please rate your experience in using the desktop root analysis software NMRooting *

❶ Choose one of the following answers
Please choose **only one** of the following:

- ◯ No experience
- ◯ Used at least once
- ◯ Sporadic user
- ◯ Expert

#### Please rate your experience in using 3D software. This includes any software or video game that displays a 3D virtual environment. *

❶ Choose one of the following answers
Please choose **only one** of the following:

- ◯ No experience
- ◯ Used at least once
- ◯ Sporadic user
- ◯ Expert

#### How much experience would you say you have in using Virtual Reality (VR) systems? *

❶ Choose one of the following answers
Please choose **only one** of the following:

- ◯ No Experience
- ◯ Used at least once
- ◯ Sporadic user
- ◯ Expert

#### Tracing in Virtual Reality

This section will give you a brief introduction to the VR Root Tracing System.

### Virtual Reality Root Tracing

The VR software you are going to be using requires you to wear a *head-mounted display*. This is a device that contains two screens, producing images for each eye. With this, we can render the 3D scene twice, making it appear to your eyes as if you were looking at a 3D space. You will also be given two controllers. There is a controller for your dominant hand and for your non-dominant hand.

The controller for your dominant hand is for **tracing and selecting**, while the other controller is for **grabbing and clicking**.

You will have some time to get used to the interaction before we begin.

| **Non-dominant Hand** | **Dominant Hand** |
| --- | --- |

| Non-dominant Hand | Dominant Hand |
| --- | --- |
| 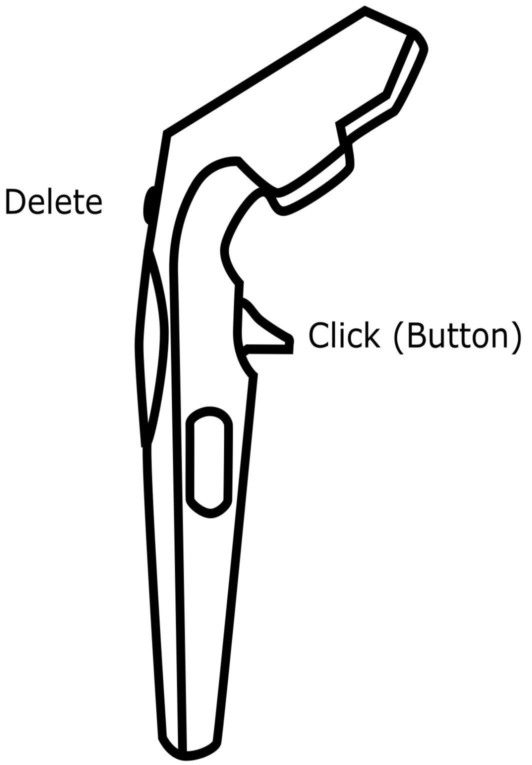  | 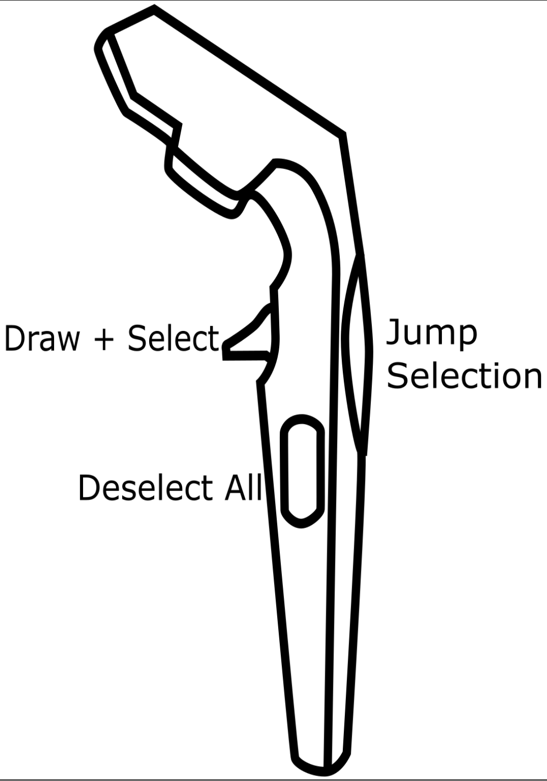  |

### The task

You will be given an isosurface volume to trace. The isosurface volume is nothing else than a computed surface that represents the volume of space occupied by matter that has a higher signal than the one chosen.

You will be asked to place a root system, that you are tracing, inside of the isosurface. What this means is that you will be placing **nodes into the geometry**. These nodes should represent the root growth inside that volume. You are allowed, and encouraged, to make educated guesses. The root surface might look a little bumpy to begin with:

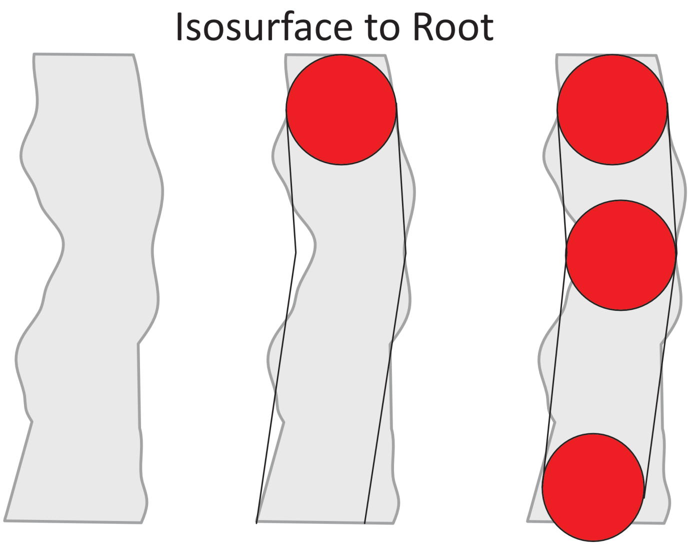


Here you will be asked to try to come up with a good "explanation" for the isosurface that might have been caused by a root. Be aware that the bumpy surface might also just be because of particular matter or local water fluxes.

### The application

You will be shown a root system MRI scan that will look something like this:

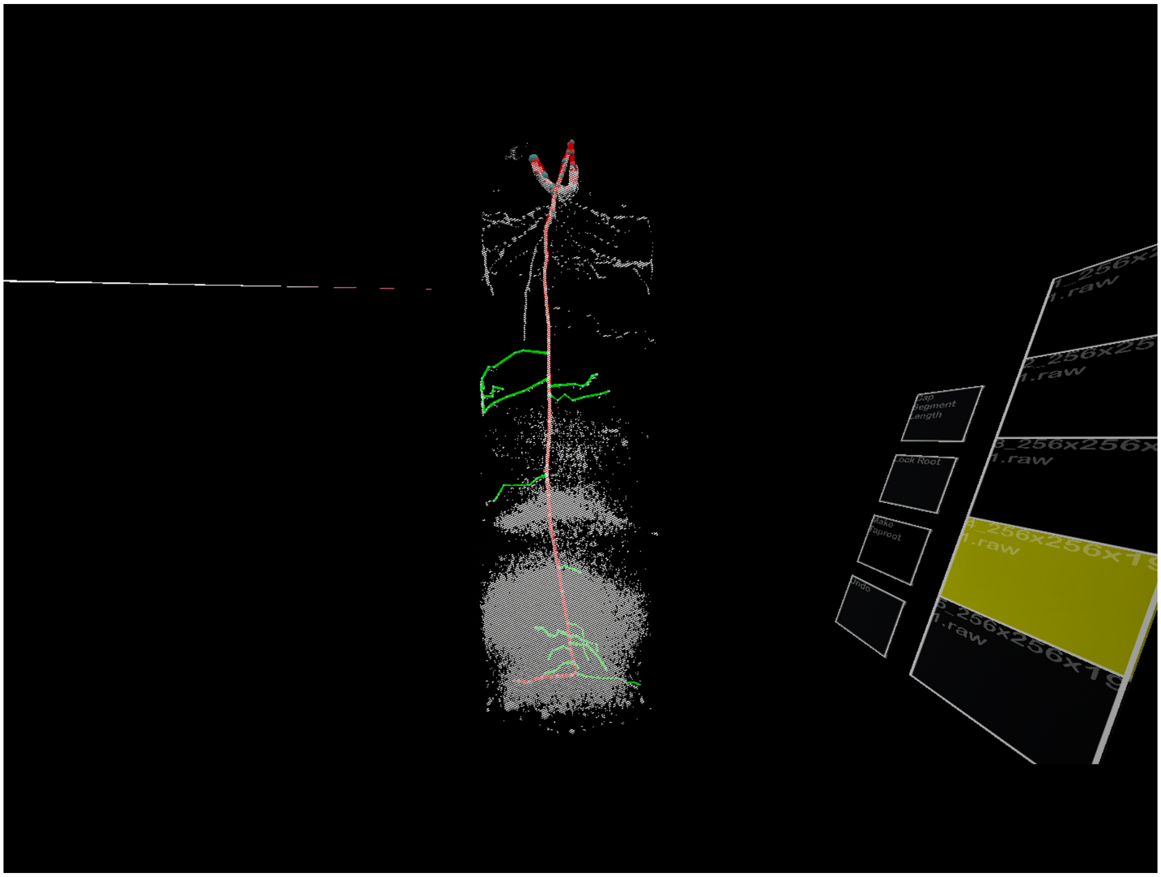

There are three components of the applications to keep track of: The **MRI scan, the root system, and the interaction widgets**.

The MRI scan is an isosurface visualization, which means that a certain signal strength is used as the surface of the root system you can see. It is semi transparent and your task is to draw a root system that resembles the MRI scan as closely as you can manage. Note that you can change the isosurface value by using the slider widget next to the menu. This slider allows you to change the signal surface for the isosurface.

You can interact with it by targeting it with the **pointing ray** and clicking with the respective trigger button (index finger).

### Please ask the supervisor about how to use the application before proceeding.

#### Indicate whether you understand to proceed *

Please choose **only one** of the following:

- ◯ Yes
- ◯ No

#### Tracing with the Desktop Application

### Annotation with the Desktop Application

We will have prepared a desktop application for you to trace the roots in. This application is called *NMRooting* and is developed by plant scientists to assist in the annotation of MRI and NMR scans.

The user interface of NMRooting looks like this:

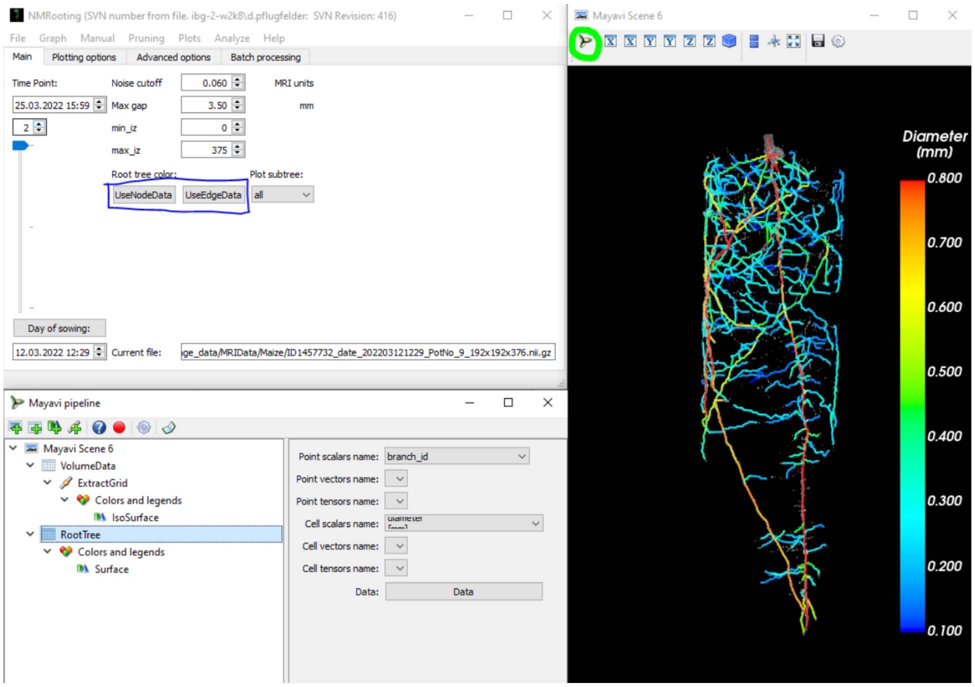


### Annotation

The data will already be ready when you change to that application. From here, you have three annotation modes. For this study, we would like you to focus on the connection of two points using NMRooting.

To do that, you select a voxel where you believe a root is starting. Afterwards, you select a voxel where the root ends.

### Interaction

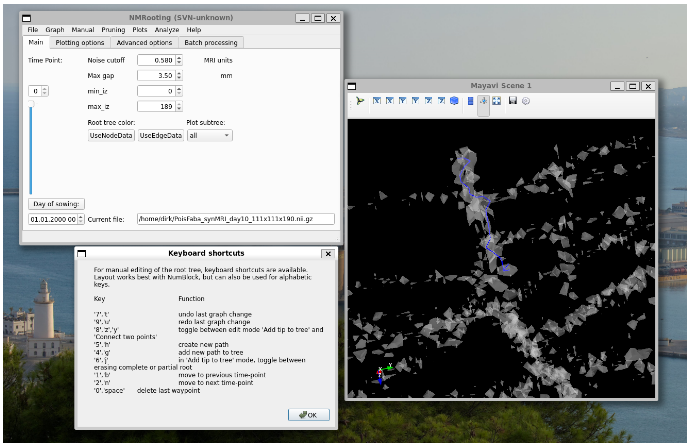


The two functions we would like you to focus on for the duration of the study are tip-to-tree and connect-two-points.

To connect a tip to the current tree, click on the data at the point where you would like the tip to be added.

The algorithm will choose the pathing to the root that you have created so far.

Note that "tree" in this instance refers to the connected voxels that are thus far labeled as part of the root.

### Root Order Labeling

Last step of the process is the labeling of the root order. For this step, you open the menu "Graph" and click on "Set Root Class".

Then, a window will open with which you can adapt what your click means in the scene. Set to "primary root", clicking on the taproot will label it as such.

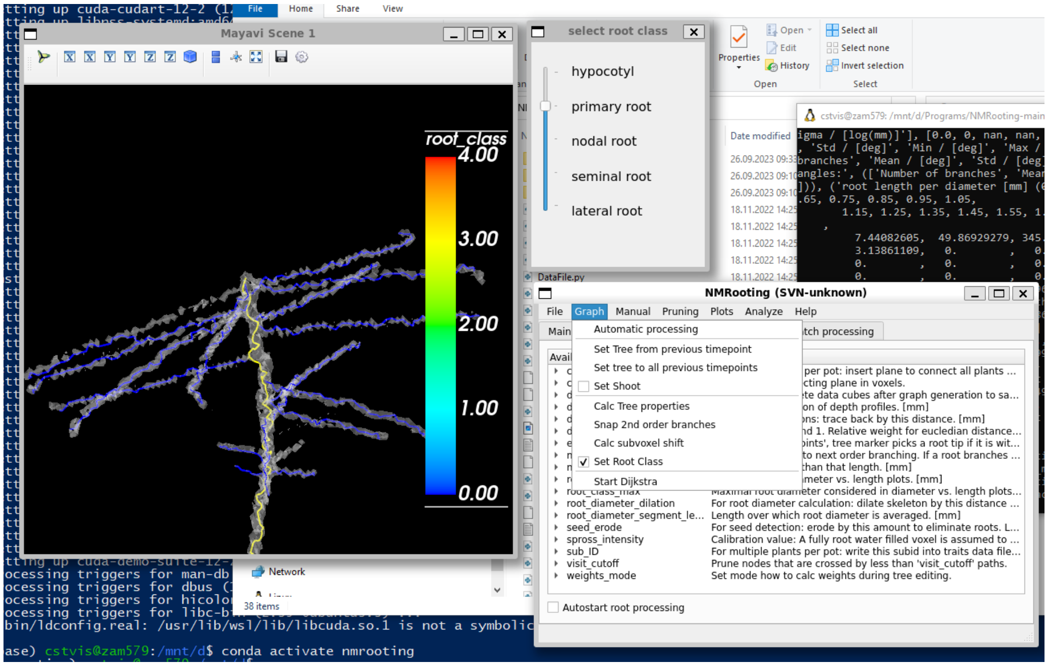


##### Indicate whether you understand to proceed *

Please choose **only one** of the following:

- ◯ Yes
- ◯ No

#### System Usability

##### I think that I would like to use this system frequently. *

❶ Choose one of the following answers

Please choose **only one** of the following:

- ◯ Strongly Disagree
- ◯ Somewhat Disagree
- ◯ Neither Disagree nor Agree
- ◯ Somewhat Agree
- ◯ Strongly Agree

#### I found the system unnecessarily complex. *

Choose one of the following answers
Please choose **only one** of the following:

- Strongly Disagree
- Somewhat Disagree
- Neither Disagree nor Agree
- Somewhat Agree
- Strongly Agree

#### I thought the system was easy to use. *

Choose one of the following answers
Please choose **only one** of the following:

- Strongly Disagree
- Somewhat Disagree
- Neither Disagree nor Agree
- Somewhat Agree
- Strongly Agree

#### I think that I would need the support of a technical person to be able to use this system. *

Choose one of the following answers
Please choose **only one** of the following:

- Strongly Disagree
- Somewhat Disagree
- Neither Disagree nor Agree
- Somewhat Agree
- Strongly Agree

#### I found the various functions in this system were well integrated. *

❗ Choose one of the following answers
Please choose **only one** of the following:

- ◯ Strongly Disagree
- ◯ Somewhat Disagree
- ◯ Neither Disagree nor Agree
- ◯ Somewhat Agree
- ◯ Strongly Agree

#### I thought there was too much inconsistency in this system. *

❗ Choose one of the following answers
Please choose **only one** of the following:

- ◯ Strongly Disagree
- ◯ Somewhat Disagree
- ◯ Neither Disagree nor Agree
- ◯ Somewhat Agree
- ◯ Strongly Agree

#### I would imagine that most people would learn to use this system very quickly. *

❗ Choose one of the following answers
Please choose **only one** of the following:

- ◯ Strongly Disagree
- ◯ Somewhat Disagree
- ◯ Neither Disagree nor Agree
- ◯ Somewhat Agree
- ◯ Strongly Agree

#### I found the system very cumbersome to use. *

❗ Choose one of the following answers
Please choose **only one** of the following:

- ◯ Strongly Disagree
- ◯ Somewhat Disagree
- ◯ Neither Disagree nor Agree
- ◯ Somewhat Agree
- ◯ Strongly Agree

I felt very confident using the system. *

Choose one of the following answers
Please choose **only one** of the following:

- Strongly Disagree
- Somewhat Disagree
- Neither Disagree nor Agree
- Somewhat Agree
- Strongly Agree

I needed to learn a lot of things before I could get going with this system. *

Choose one of the following answers
Please choose **only one** of the following:

- Strongly Disagree
- Somewhat Disagree
- Neither Disagree nor Agree
- Somewhat Agree
- Strongly Agree

### User Experience

For the assessment of the application, please fill out the following questionnaire. The questionnaire consists of pairs of contrasting attributes that may apply to the product. The circles between the attributes represent gradations between the opposites. You can express your agreement with the attributes by ticking the circle that most closely reflects your impression.

### Example:

attractive ○ ○ ○ ○ ○ ○ ○ unattractive

This response would mean that you rate the application as more attractive than unattractive.

Please decide spontaneously. Don’t think too long about your decision to make sure that you convey your original impression.

Sometimes you may not be completely sure about your agreement with a particular attribute or you may find that the attribute does not apply completely to the particular product. Nevertheless, please tick a circle in every line.

It is your personal opinion that counts. Please remember: there is no wrong or right answer!

#### Please assess the application now by ticking one circle per line. *

Please choose the appropriate response for each item:

| | 1 | 2 | 3 | 4 | 5 | 6 | 7 | |
|---|---|---|---|---|---|---|---|---|
| **annoying** | | | | | | | | **en** |
| **not understandable** | | | | | | | | **un** |
| **creative** | | | | | | | | **du** |
| **easy to learn** | | | | | | | | **dif to lea** |
| **valuable** | | | | | | | | **inf** |
| **boring** | | | | | | | | **ex** |
| **not interesting** | | | | | | | | **int** |
| **unpredictable** | | | | | | | | **pr** |
| **fast** | | | | | | | | **sl** |
| **inventive** | | | | | | | | **co** |
| **obstructive** | | | | | | | | **su** |
| **good** | | | | | | | | **ba** |
| **complicated** | | | | | | | | **ea** |
| **unlikable** | | | | | | | | **pl** |
| **usual** | | | | | | | | **lea ed** |
| **unpleasant** | | | | | | | | **pl** |
| **secure** | | | | | | | | **no se** |
| **motivating** | | | | | | | | **de** |
| **meets expectations** | | | | | | | | **do no me ex** |
| **inefficient** | | | | | | | | **eff** |
| **clear** | | | | | | | | **co** |

| | 1 | 2 | 3 | 4 | 5 | 6 | 7 | |
|---|---|---|---|---|---|---|---|---|
| **impractical** | | | | | | | | **pr** |
| **organized** | | | | | | | | **cl** |
| **attractive** | | | | | | | | **un** |
| **friendly** | | | | | | | | **un** |
| **conservative** | | | | | | | | **in** |

### NASA-TLX Short

Please rate the application in the 10-point diverging scale below.

#### Physical Demand

How physically demanding was the task?

*

Please choose the appropriate response for each item:

| | | | | | | | | | | | | | | | | | | | | | |
|---|---|---|---|---|---|---|---|---|---|---|---|---|---|---|---|---|---|---|---|---|---|
| **Very Low** | | | | | | | | | | | | | | | | | | | | | **Ve Hi** |

#### Temporal Demand

How hurried or rushed was the pace of the task?

*

Please choose the appropriate response for each item:

| | | | | | | | | | | | | | | | | | | | | | |
|---|---|---|---|---|---|---|---|---|---|---|---|---|---|---|---|---|---|---|---|---|---|
| **Very Low** | | | | | | | | | | | | | | | | | | | | | **Ve Hi** |

#### Performance

How successful were you in accomplishing what you were asked to do?

*

Please choose the appropriate response for each item:

| | | | | | | | | | | | | | | | | | | | | | |
|---|---|---|---|---|---|---|---|---|---|---|---|---|---|---|---|---|---|---|---|---|---|
| **Very Low** | ○ | ○ | ○ | ○ | ○ | ○ | ○ | ○ | ○ | ○ | ○ | ○ | ○ | ○ | ○ | ○ | ○ | ○ | ○ | ○ | **Ve Hi** |

#### Effort

How hard did you have to work to accomplish your level of performance?

*

Please choose the appropriate response for each item:

| | | | | | | | | | | | | | | | | | | | | | |
|---|---|---|---|---|---|---|---|---|---|---|---|---|---|---|---|---|---|---|---|---|---|
| **Very Low** | ○ | ○ | ○ | ○ | ○ | ○ | ○ | ○ | ○ | ○ | ○ | ○ | ○ | ○ | ○ | ○ | ○ | ○ | ○ | ○ | **Ve Hi** |

#### Frustration

How insecure, discouraged, irritated, stressed, and annoyed were you?

*

Please choose the appropriate response for each item:

| | | | | | | | | | | | | | | | | | | | | | |
|---|---|---|---|---|---|---|---|---|---|---|---|---|---|---|---|---|---|---|---|---|---|
| **Very Low** | ○ | ○ | ○ | ○ | ○ | ○ | ○ | ○ | ○ | ○ | ○ | ○ | ○ | ○ | ○ | ○ | ○ | ○ | ○ | ○ | **Ve Hi** |

#### Tracing in Virtual Reality

This section will give you a brief introduction to the VR Root Tracing System.

### Virtual Reality Root Tracing

The VR software you are going to be using requires you to wear a *head-mounted display*. This is a device that contains two screens, producing images for each eye. With this, we can render the 3D scene twice, making it appear to your eyes as if you were looking at a 3D space. You will also be given two controllers. There is a controller for your dominant hand and for your non-dominant hand.

The controller for your dominant hand is for **tracing and selecting**, while the other controller is for **grabbing and clicking**.

You will have some time to get used to the interaction before we begin.

| **Non-dominant Hand** | **Dominant Hand** |
|---|---|
